# Selective activation of Gαob by an adenosine A1 receptor agonist elicits analgesia without cardiorespiratory depression

**DOI:** 10.1101/2020.04.04.023945

**Authors:** Mark J. Wall, Emily Hill, Robert Huckstepp, Kerry Barkan, Giuseppe Deganutti, Michele Leuenberger, Barbara Preti, Ian Winfield, Sabrina Carvalho, Anna Suchankova, Haifeng Wei, Dewi Safitri, Xianglin Huang, Wendy Imlach, Circe La Mache, Eve Dean, Cherise Hume, Stephanie Hayward, Jess Oliver, Fei-Yue Zhao, David Spanswick, Christopher A. Reynolds, Martin Lochner, Graham Ladds, Bruno G. Frenguelli

## Abstract

The development of therapeutic agonists for G protein-coupled receptors (GPCRs) is hampered by the propensity of GPCRs to couple to multiple intracellular signalling pathways. This promiscuous coupling leads to numerous downstream cellular effects, some of which are therapeutically undesirable. This is especially the case for adenosine A1 receptors (A_1_Rs) whose clinical potential is undermined by the sedation and cardiorespiratory depression caused by conventional agonists. We have discovered that the A_1_R-selective agonist, BnOCPA, is a potent and powerful analgesic but does not cause sedation, bradycardia, hypotension or respiratory depression. This unprecedented discrimination between native A_1_Rs arises from BnOCPA’s unique and exquisitely selective activation of Gob among the six Gαi/o subtypes, and in the absence of β-arrestin recruitment. BnOCPA thus demonstrates a highly-specific Gα-selective activation of the native A_1_R, sheds new light on GPCR signalling, and reveals new possibilities for the development of novel therapeutics based on the far-reaching concept of selective Gα agonism.

**Short summary:** We describe the selective activation of an adenosine A1 receptor-mediated intracellular pathway that provides potent analgesia in the absence of sedation or cardiorespiratory depression, paving the way for novel medicines based on the far-reaching concept of selective Gα agonism.

## Introduction

G protein-coupled receptors (GPCRs) are the targets of many FDA-approved drugs^1, 2^. However, the promiscuity with which they couple to multiple G protein- and β-arrestin-activated intracellular signalling cascades leads to unwanted side effects. These side effects limit both the range of GPCRs suitable for drug-targeting, and the number of conditions for which treatments could be developed^3, 4^. One family of GPCRs that has particularly suffered as drug targets from their promiscuous coupling and wide-ranging cellular actions are the four GPCRs for the purine nucleoside adenosine, despite the potential for using adenosine receptor agonists to treat many pathological conditions including cancer, and various cardiovascular, neurological and inflammatory diseases^5–7^. For example, activation of the widely-distributed adenosine A1 receptor (A_1_R) with currently available agonists elicits multiple actions in both the central nervous system (CNS) and in the cardiorespiratory system. In the CNS A_1_Rs inhibit synaptic transmission, induce neuronal hyperpolarization, and cause sedation, while in the cardiorespiratory system A_1_Rs slow the heart (bradycardia), contribute to reducing blood pressure (hypotension), and depress respiration (dyspnea)^7–12^. These multiple effects severely limit the prospects of A_1_R agonists as life-changing medicines, despite their potential use in a wide range of clinical conditions, such as glaucoma, type 2 diabetes mellitus, pain, epilepsy and cerebral ischemia^7, 13–16^, and in which there are clear unmet clinical needs that could be addressed with novel therapeutics.

The therapeutic limitations of promiscuous GPCR coupling might be overcome through the development of biased agonists – compounds that selectively recruit one intracellular signalling cascade over another^4, 17, 18^. This signalling bias has most frequently been expressed in terms of Gα vs β-arrestin signalling^19^ and has been pursued at a variety of receptors^20, 21^, for example, at the angiotensin II type 1 receptor (AT1R)^22^, and at neurotensin receptors in the treatment of drug addiction^23^. Agonist bias has been sought in the context of opioid receptors, but with some controversy^24^, for compounds producing analgesia with reduced respiratory depression^4^.

However, while other forms of bias exist, including between individual Gα subunits^17, 25, 26^, the challenge remains in translating GPCR signalling bias observed *in vitro* to tangible, and physiologically- and clinically-relevant, selectivity at native receptors *in vivo*^3, 4, 27, 28^. Accordingly, while the potential to preferentially drive the G protein-coupling of A_1_Rs has been described in several *in vitro* studies^29–32^, to date no A_1_R-specific agonist has been reported that can elicit biased Gα agonism at native A_1_Rs in intact physiological systems, let alone the selective activation of one Gα subunit. To achieve such selectivity among Gα subunits would introduce novel therapeutic opportunities across a wide range of debilitating clinical conditions.

Utilising molecular dynamics (MD) simulations, and Gαi/o subunit- and β-arrestin-specific cellular signalling assays, we describe how one A_1_R-selective agonist, BnOCPA^33, 34^, fulfils the criteria for a selective Gα agonist in exclusively activating Gob among the six members of the Gαi/o family of G protein subunits, and in the absence of β-arrestin recruitment. In addition, through a combination of CNS electrophysiology, physiological recordings of cardiorespiratory parameters, a sensitive assay of attention and locomotor function, and the use of a clinically-relevant model of chronic neuropathic pain, we demonstrate selective activation of native A_1_Rs and the delivery of potent analgesia without sedation, motor impairment or cardiorespiratory depression. Our data thus demonstrate the translation of agonist Gα selectivity *in vitro* to therapeutically-tangible clinically-relevant observations *in vivo*. Such observations reveal the possibility of achieving Gα selectivity at native receptors, highlight the physiological benefits of such selectivity, and specifically speak to the possibility of unlocking the widespread clinical potential of A_1_R agonists.

## RESULTS

### The novel A_1_R agonist BnOCPA exquisitely discriminates between native pre- and postsynaptic A_1_Rs in the intact mammalian CNS

BnOCPA (Fig. 1a), a molecule first described in a patent as a potential treatment for glaucoma or ocular hypertension^34^, is a cyclopentyl derivative of adenosine and a highly selective and potent, full agonist at human adenosine A_1_Rs (hA_1_Rs; Fig. 1b; Supplementary Table 1)^33^. Our characterisation of BnOCPA, synthesised independently as part of a screen for suitable scaffolds for the generation of fluorescent ligands for the A_1_R, began with an exploration of the binding characteristics of BnOCPA at the human A_1_R (hA_1_R) using classical radioligand binding (where the antagonist [^3^H]DPCPX was used as a tracer), and a NanoBRET agonist binding assay (using a novel NECA-TAMRA compound, which acts as a full agonist (pEC_50_ – 7.23 ± 0.13; *See Methods*). Using both assays we observed that BnOCPA was able to bind to the hA_1_R with an affinity equal to that of the prototypical A_1_R agonists CPA and NECA, and higher than that of the endogenous agonist adenosine (Fig. 1b; Supplementary Table 1). Significantly, using NECA-TAMRA as the fluorescent agonist tracer, the high affinity state of the biphasic binding profile observed in the NanoBRET assay was equivalent to that reported previously for BnOCPA (3.8 nM compared to 1.7 nM^34^).

**Fig. 1.**
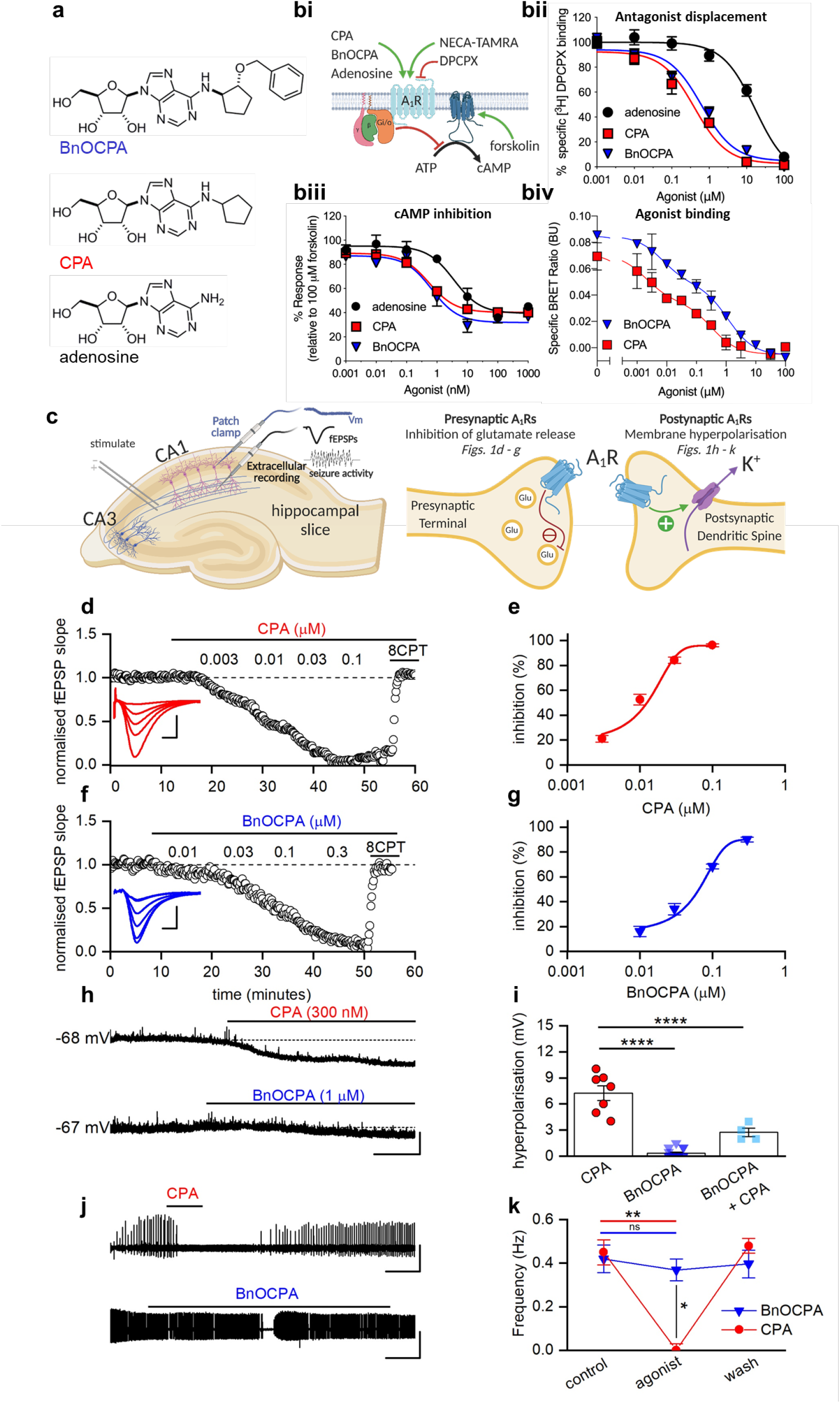
BnOCPA discriminates between pre- and postsynaptic A_1_Rs in the CNS. **a** Chemical structures of adenosine, CPA and its derivative, BnOCPA^33^. **bi:** Schematic of the binding of adenosine, CPA and BnOCPA to the human (h) A_1_R was measured via their ability to displace [^3^H]DPCPX, a selective antagonist for the A_1_R, from membranes prepared from CHO-K1-hA_1_R cells, their ability to elicit Gi/o-mediated inhibition of forskolin-stimulated production of cAMP, and for CPA and BnOCPA to displace binding of the fluorescent AR agonist NECA-TAMRA in a BRET assay. **bii:** CPA and BnOCPA bind with equal affinity to the A_1_R (pK_i_ ∼6.6), while adenosine has a reduced affinity (pK_i_ ∼5; *n* = 5 – 19 individual repeats). **biii:** cAMP levels were measured in CHO-K1-hA_1_R cells following co-stimulation with 1 μM forskolin and each compound (1 pM – 1 μM) for 30 minutes. This identified that all are full agonists at the hA_1_R. Adenosine displayed a 10-fold reduced potency compared to CPA and BnOCPA (*n* = 4 individual repeats). **biv**: CPA and BnOCPA displace the fluorescent AR agonist NECA-TAMRA in a BRET assay revealing a bi-phasic binding profile indicating that both compounds display high affinity and low affinity binding. The high affinity constants for CPA and BnOCPA at the A_1_R were pK_i_ ∼9.02 and ∼8.44, respectively (n= 3 individual repeats) with the low affinity constants matching that stated in **bii. c** Diagram illustrating ***Left,*** hippocampal slice preparation showing position of stimulating, patch clamp and extracellular recording electrodes together with representative electrophysiological recordings: membrane potential (Vm), a fEPSP (field excitatory postsynaptic potential), which is a product of the electrical stimulation-induced release of glutamate and the activation of postsynaptic glutamate receptors (not shown), and seizure activity caused by overactivation of glutamate receptors; ***Right,*** pre- and postsynaptic A_1_Rs at hippocampal synapses, their physiological effects upon activation, and the panels in Fig. 1 where these effects can be seen (presynaptic: panels **d – g**; postsynaptic: panels **h – k**). **d** Increasing concentrations of the A_1_R agonist CPA reduced the fEPSP, an effect reversed by the A_1_R antagonist 8CPT (2 µM). The graph plots the normalised negative slope of the fEPSP, an index of the strength of synaptic transmission, over time. Inset, superimposed fEPSP averages in control (largest fEPSP) and becoming smaller in increasing concentrations of CPA. Scale bars measure 0.2 mV and 5 ms. **e** Concentration-response curve for the inhibition of synaptic transmission by CPA (IC_50_ = 11.8 ± 2.7 nM; *n* = 11 slices). **f** Increasing concentrations of BnOCPA reduced the fEPSP, an effect reversed by 8CPT (2 µM). Inset, superimposed fEPSP averages in control and in increasing concentrations of BnOCPA. Scale bars measure 0.1 mV and 2 ms. **g** Concentration-response curve for the inhibition of synaptic transmission by BnOCPA (IC_50_ = 65 ± 0.3 nM; *n* = 11 slices). **h** CPA (300 nM) hyperpolarised the membrane potential while BnOCPA (1 µM) had little effect. Scale bars measure 4 mV and 30 s. **i** Summary data for membrane potential changes. The mean hyperpolarisation produced by CPA (300 nM; 7.26 ± 0.86 mV, *n* = 7 cells) was significantly different (one-way ANOVA; F(2,23) = 70.46; P = 1.55 x 10^-10^) from that produced by BnOCPA (300 nM–-1 µM; 0.33 ± 0.14 mV, *n* = 10 and 5 cells, respectively; P = 8.26 x 10^-11^) and for CPA (300 nM) applied in the presence of BnOCPA (300 nM; 2.75 ± 0.48 mV, n = 4 cells, P = 2.89 x 10^-5^; See Supplementary Fig. 2a for an example trace). **j** In an *in vitro* model of seizure activity, represented as frequent spontaneous spiking from baseline, CPA (300 nM) reversibly blocked activity while BnOCPA (300 nM) had little effect. Scale bars measure 0.5 mV and 200 s. **k** Summary data for seizure activity expressed in terms of the frequency of spontaneous spiking before, during and after CPA or BnOCPA. CPA abolished seizure activity (*n* = 4) whereas BnOCPA did not significantly reduce seizure frequency (*n* = 6). Data represented as mean ± SEM; Two-way RM ANOVA (BnOCPA vs CPA slices): F(1, 3) = 186.11, P = 8.52 x10^-4^ with the following Bonferroni post hoc comparisons: BnOCPA vs Control; P = 1; CPA vs control; P = 0.010; BnOCPA vs CPA; P = 0.027. Averaged data is presented as mean ± SEM. ns, not significant; *, P < 0.05; **, P < 0.02; ****, P < 0.0001.

These initial pharmacological studies at recombinant hA_1_Rs in cell lines did not reveal anything extraordinary about BnOCPA. However, when we investigated BnOCPA at native A_1_Rs in rat hippocampal slices, against which BnOCPA is also a potent agonist, with ∼8,000- and >150-fold greater efficacy at rat A_1_Rs (rA_1_Rs) than at rat A_2_ARs (rA_2_ARs) and A_3_Rs (rA_3_Rs), respectively (Supplementary Table 2), we discovered properties of BnOCPA that were not consistent with those of typical A_1_R agonists such as adenosine, CPA and NECA. In accordance with the effects of standard A_1_R agonists, BnOCPA potently inhibited excitatory synaptic transmission in rat hippocampal slices (IC_50_ ∼65 nM; Fig. 1c to g and Supplementary Fig.1a to d). This effect was attributable to activation of native presynaptic A_1_Rs on glutamatergic terminals^9^ (Fig. 1c; Supplementary Fig. 1e, f), and cannot be attributed to any action of BnOCPA at A_3_Rs since even a high concentration (1 µM) of the potent and selective A_3_R agonist 2-Cl-IB-MECA^35^ had no effect on synaptic transmission (Supplementary Fig. 1g, h). However, in stark contrast to adenosine and CPA, BnOCPA did not activate postsynaptic A_1_Rs (Fig. 1c) to induce membrane hyperpolarisation, even at concentrations 15 times the IC_50_ for the inhibition of synaptic transmission (Fig. 1h, i).

This peculiar and unique discrimination between pre- and postsynaptic A_1_Rs might possibly be explained in terms of either some hindrance in the binding of BnOCPA to A_1_Rs on postsynaptic neurones, or, and unprecedented for an A_1_R agonist, binding to the postsynaptic A_1_R, but without the ability to activate the receptor. To test the latter hypothesis – that BnOCPA actually bound to postsynaptic A_1_Rs, but without efficacy we reasoned that BnOCPA might behave in a manner analogous to a receptor antagonist in preventing or reversing activation by other A_1_R agonists, a property that has been predicted and observed for biased agonists at other receptors^17, 27^. To test this, we pre-applied BnOCPA then applied CPA (in the continued presence of BnOCPA). Remarkably, the co-application of CPA and BnOCPA resulted in a significant reduction of the effects of CPA on membrane potential (Fig. 1i; Supplementary Fig. 2a, b). In addition, membrane hyperpolarisation induced by the endogenous agonist adenosine was reversed by BnOCPA (Supplementary Fig. 2c). In contrast, the A_3_R agonist 2-Cl-IB-MECA had no effect on membrane potential and did not interfere with the membrane hyperpolarisation caused by adenosine (Supplementary Fig. 2d, e), further reaffirming the actions of BnOCPA as being selectively mediated by A_1_Rs.

To test whether the inability of BnOCPA to affect membrane potential was a trivial action due to BnOCPA blocking K^+^ channels mediating the postsynaptic hyperpolarisation, or in some other way non-specifically interfering with G protein signalling, we applied the GABA_B_ receptor agonist baclofen to CA1 pyramidal neurons. BnOCPA had no effect on membrane hyperpolarisation produced by baclofen (Supplementary Fig. 2f, g), confirming that the actions of BnOCPA were specific to the A_1_R. These observations, of a lack of effect of BnOCPA on postsynaptic membrane potential, likely explained why, in a model of seizure activity, (low Mg^2+^/high K^+^), with prominent postsynaptic depolarisation that promotes neuronal firing, BnOCPA had little effect (Fig. 1j, k). In contrast, equivalent concentrations of CPA completely suppressed neuronal firing (Fig. 1jw, k).

### BnOCPA demonstrates unique Gα signalling in the selective activation of Gob

The observation that BnOCPA discriminated between pre- and postsynaptic A_1_Rs might be explained if these receptors were to activate different intracellular pathways to mediate their effects, and that BnOCPA was not able to activate the pathway responsible for postsynaptic membrane hyperpolarisation. To test whether the actions of BnOCPA and the prototypical A_1_R agonists were mediated via β-arrestins (β-arrestin1 and β-arrestin2), we used a BRET assay^36–40^ for β-arrestin recruitment (Supplementary Fig. 3). We observed no β-arrestin recruitment at the A_1_R using either BnOCPA, CPA or adenosine, regardless of whether β-arrestin1 or β-arrestin2 was expressed (Supplementary Fig. 3). This was in contrast to β-arrestin2 recruitment by the A_3_R in response to adenosine and NECA, but not BnOCPA (Supplementary Fig. 3). Moreover, the lack of recruitment of β-arrestin1 and β-arrestin2 by the A_1_R was independent of any of the six G protein receptor kinase (GRK) isoforms co-expressed with β-arrestin1 and β-arrestin2; only low levels of recruitment were observed even when GRKs where highly (5-fold) over-expressed compared to the levels in the A_3_R assays (Supplementary Fig. 4). These observations of a lack of β-arrestin recruitment by A_1_Rs are consistent with those previously reported for recombinant A_1_Rs expressing native sequences^41–45^, and are likely due to the absence of serine and threonine residues in the A_1_R cytoplasmic tail, which makes the A_1_R intrinsically biased against β-arrestin signalling^19, 46^. Accordingly, the differential actions of BnOCPA at pre- and postsynaptic A_1_Rs are more likely to reside in selective activation of one Gα-mediated pathway over another.

To investigate whether BnOCPA has the ability to discriminate between the various Gαi/o subunits activated by adenosine, we generated a recombinant cell system (CHO-K1 cells) expressing both the hA_1_R and individual pertussis toxin (PTX)-insensitive variants of individual Gαi/o subunits. Against these individual Gαi/o subunits we tested adenosine, CPA, NECA, BnOCPA, and the agonist HOCPA^33, 47^, a stereoisomer of GR 79236^48, 49^, which behaved similarly to adenosine and CPA in both inhibiting synaptic transmission and causing membrane hyperpolarisation (Supplementary Fig. 5). In cells treated with PTX to inhibit endogenous Gαi/o^30, 33^ we observed that adenosine, CPA, NECA and HOCPA activated a range of Gα subunits. Common to all of these agonists was the activation of both Gαo isoforms, Goa and Gob, with differential activation of Gi1 (HOCPA), Gi2 (NECA, CPA) and Gz (adenosine; Fig. 2a-e; Supplementary Figs. 5 and 6). Such promiscuous and biased Gα coupling has been described previously for adenosine, CPA, and NECA at recombinant A_1_Rs in cell lines^29, 50^, including using novel BRET-based assays for adenosine at some Gαi/o^51^. These previous observations are in keeping with ours, confirming the validity of the PTX-based approach. In stark contrast, BnOCPA displayed a unique and highly distinctive Gαi/o subunit activation profile: BnOCPA was not able to activate Gi1, Gi2, Gi3 or Gz, and was furthermore capable of discriminating between the two Gαo isoforms via the selective activation of Gob, and not of Goa (Fig. 2a-e; Supplementary Fig. 6).

**Fig. 2.**
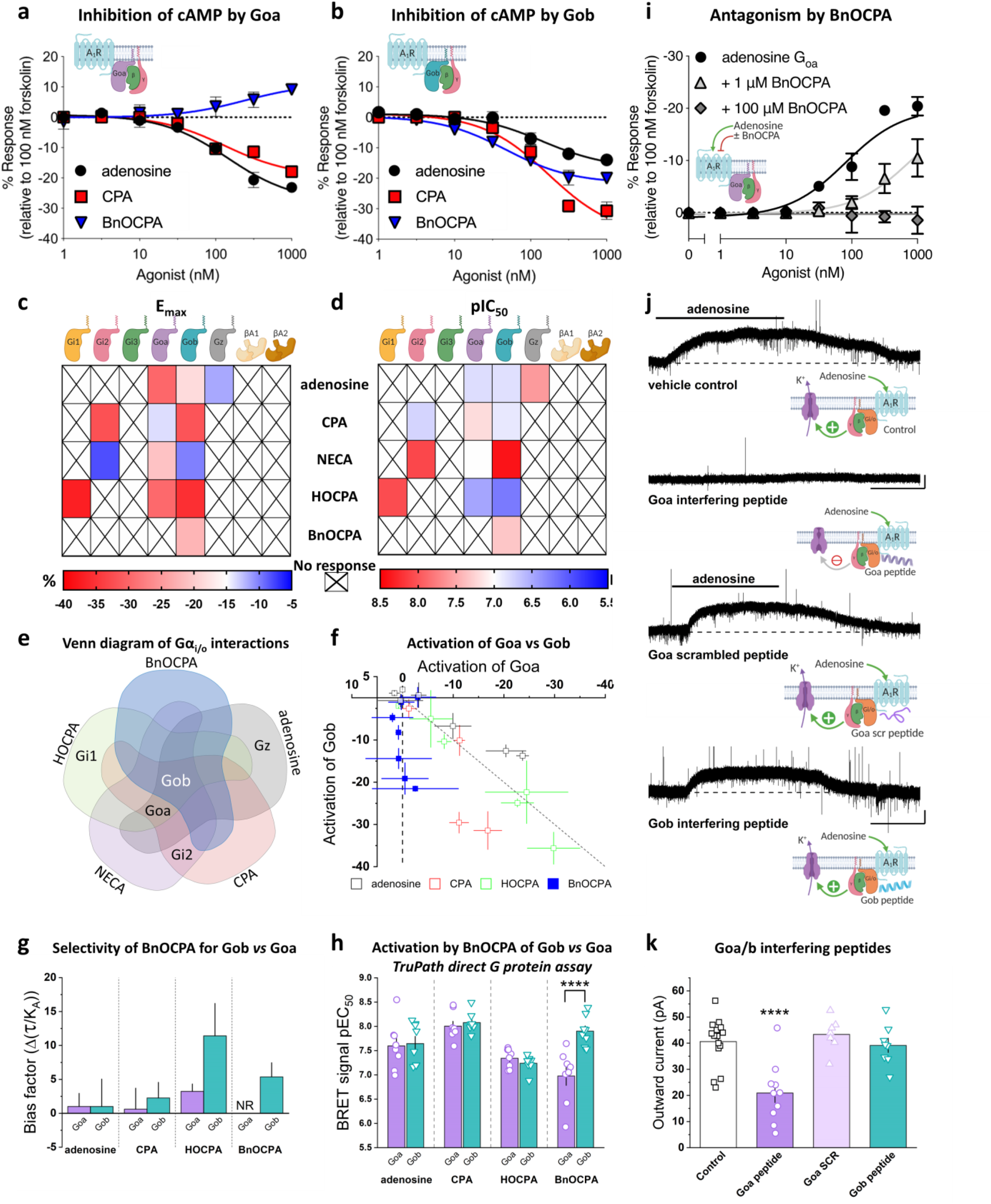
BnOCPA selectively activates Gob. **a** cAMP accumulation was measured in PTX-pre-treated (200 ng/ml) CHO-K1-hA_1_R cells expressing PTX-insensitive Goa following co-stimulation with 1 μM forskolin and each compound (1 pM–-1 μM) for 30 minutes (*n =* 6 individual repeats). The data demonstrates that BnOCPA does not activate Goa. **b**, as for **a** but cells were transfected with PTX-insensitive Gob. Adenosine, CPA and BnOCPA all inhibit cAMP accumulation through coupling to Gob (*n* = 6 individual repeats). Stimulation of cAMP production in **a** reflects BnOCPA’s activation of endogenous, PTX-resistant Gαs by the A_1_R and is in agreement with previous observations for other A_1_R agonists (see Supplementary Figs. 5 and 6 and^29, 106, 107^). **c-d** Heatmaps summarising E_max_ (**c; %**) and potency (**d**; pEC50; -log [agonist concentration] required for 50 % inhibition of cAMP accumulation) for individual Gα subunit and β arrestin1 and 2 activation by selective A_1_R agonists for the inhibition of forskolin-stimulated cAMP production. Data taken from: adenosine, CPA, BnOCPA Fig. 1, Supplementary Figs. 3, 5, 6; NECA, Supplementary Fig. 3, 5, 6; HOCPA, Supplementary Fig. 5. **e** Venn diagram of agonist interactions with individual Gαo/i subunits. While adenosine, CPA, NECA and HOCPA each activate three subunits, BnOCPA exclusively activates one, Gob. **f** The inhibition of cAMP accumulation via A_1_R:Goa or A_1_R:Gob by the endogenous agonist adenosine, and the selective A_1_R agonists CPA, HOCPA and BnOCPA. Each data point represents a concentration of agonist from the data in Supplementary Figs. 5 and 6. Equal activation of Goa and Gob at each concentration (no bias) would fall on the line of identity (broken grey line). HOCPA behaves most like a Goa/Gob unbiased agonist, with some bias for Gob shown by CPA, and for Goa by adenosine. BnOCPA is highly selective for Gob, with no activation of Goa, as indicated by the data points falling on the line of zero Goa activation (vertical broken line). Data presented as mean ± SEM and is replotted from Supplementary Figs. 5 and 6. **G** Signalling bias of A_1_R-selective agonists for A_1_R-Goa and A_1_R-Gob (Δ(τ/K_A_)) was determined relative to the natural agonist adenosine using the change in (τ/K_A_) ratio for the data in **f** where τ is the efficacy of each agonist in activating individual A_1_R-Gɑi/o complexes, and where K_A_ is the agonist equilibrium dissociation constant. Compared to adenosine BnOCPA elicits no measurable response (NR) at Goa. **h** The TruPath assay for direct G protein activation reveals no preference between Goa and Gob by adenosine, CPA or HOCPA, but a significant >10-fold greater activation of Gob *vs* Goa by BnOCPA (unpaired Student’s t-test; P = 0.0009; see also Supplementary Fig. 7A). **i** Adenosine’s ability to inhibit cAMP accumulation via its activation of Goa was inhibited by BnOCPA in a concentration-dependent manner, and with a K_d_ of 113 nM (pK_d_ ∼6.9 (*n* = 4 individual repeats) similar to the binding affinity to the hA_1_R pK_i_ ∼6.6; Fig. 1B). No agonist action of BnOCPA is observed at Goa even at high concentrations. **j** Example current traces produced by adenosine (10 µM) in control conditions or in the presence of intracellular Goa interfering peptide (100 µM), scrambled Goa peptide (100 µM) or Gob interfering peptide (100 µM). Scale bars measure 25 pA and 100 s. **k** Summary data of adenosine-induced outward current experiments. The mean amplitude of the outward current induced by adenosine (40.6 ± 2.2 pA, *n* = 16 cells) was significantly reduced (one-way ANOVA; F(3,37) = 12.40, P = 9.22 x 10^-6^) to 20.9 ± 3.6 pA (*n* = 10 cells, P = 2.65 x 10^-5^) in 100 µM Goa interfering peptide. Neither the scrambled Goa peptide (Goa SCR; 43.4 ± 2.4 pA, *n* = 7 cells, P = 1) nor the Gob interfering peptide (39. 2 ± 2.7 pA, *n* = 8 cells, P = 1) significantly reduced the amplitude of the adenosine-induced outward current compared to control, but each were significantly different from the Goa interfering peptide (P = 8.20 x 10^-5^; P = 8.86 x 10^-4^, respectively). Averaged data is presented as mean ± SEM. ****, P < 0.0001 relative to other groups.

The selective and unique activation of Gob among the six Gαi/o subunits by BnOCPA could be observed in a comparison of the activation of Goa and Gob by the native and selective A_1_R agonists in their ability to inhibit the forskolin-stimulated accumulation of cAMP (Fig. 2f). Whereas adenosine, CPA and HOCPA activated both Goa and Gob to inhibit cAMP accumulation, BnOCPA selectively activated Gob, with no discernible activation of Goa. Further quantification of this Gα selectivity, through the application of the operational model of receptor agonism^52–54^ to remove potential issues of system bias, confirmed selective activation of Gob by BnOCPA, with no detectable response at Goa (Fig. 2g). As further validation of the ability of BnOCPA to discriminate between the activation of Goa and Gob, we took advantage of the recently described TruPath assay^55^, which utilises a reduction in a Gα-Gβγ BRET signal to infer agonist-induced G protein activation (Fig. 2h; Supplementary Fig. 7a). Adenosine, CPA, and HOCPA elicited equipotent activation of both Goa and Gob. In stark contrast to these agonists, BnOCPA was >10-fold more efficacious in activating Gob than Goa, and, of all the agonists tested, BnOCPA displayed the weakest potency at Goa. While subtle differences between the Goa and Gob response exist across the two very different *in vitro* assays, these data nonetheless confirm that BnOCPA demonstrates a previously unprecedented ability for an A_1_R agonist to discriminate between Gα subtypes.

To establish the functional implications of BnOCPA’s profound selectivity for Gob over Goa, we hypothesised that BnOCPA should reduce the actions of adenosine on the inhibition of cAMP accumulation via Goa. This was indeed the case (Fig. 2i): BnOCPA antagonised the Goa-mediated inhibition of cAMP production by adenosine in a concentration-dependent manner. This classic attribute of an antagonist enabled a Schild analysis estimate of BnOCPA’s affinity (Kd) to be 113 nM, with a pKd ∼ 6.9^56^, a value that was quantitatively similar to BnOCPA’s ability to bind to the hA_1_R (pK_i_ ∼6.6; Fig. 1b). Importantly, this observation, of the ability of BnOCPA to antagonise the actions of adenosine on cAMP inhibition (Fig. 2i), revealed no agonist action of BnOCPA at Goa at concentrations up to 100 μM (>10^5^ greater than the IC_50_ against cAMP accumulation; Fig. 1b and ∼10^4^ greater than the EC50 in the TruPath assay; Fig. 2h), and, moreover, had parallels with the antagonising effects of BnOCPA on membrane potential in the CNS (Fig. 1h, i; Supplementary Fig. 2a, c). These data suggest that BnOCPA has the unique ability of displaying both agonist and antagonist-like properties at both recombinant and native A_1_Rs; properties that are expected of a truly Gα subunit-selective agonist.

The data from whole-cell patch-clamp recordings showed that BnOCPA did not influence neuronal membrane potential at native A_1_Rs (Fig. 1h, i), while experiments in recombinant hA_1_Rs showed that BnOCPA did not activate Goa (Fig. 2a, c-f), and indeed *prevented* the activation of Goa by adenosine (Fig. 2i). We thus predicted that A_1_Rs in the hippocampus, where Goa is found at levels 10-15 times higher than Gob^57^, should act via Goa to induce membrane hyperpolarisation, and thereby providing a potential explanation for the lack of effect of BnOCPA on membrane potential. To test this prediction, we injected a series of previously-validated interfering peptides against Goa and Gob ^58–67^ into CA1 pyramidal cells during whole-cell voltage clamp recordings. Introduction of the Goa interfering peptide caused a significant attenuation of the adenosine-induced outward current (Fig. 2j, k). In contrast, neither the scrambled Goa peptide, nor the Gob peptide, which reduced the modulation of Ca^2+^ channels by muscarinic M4 receptors in striatal cholinergic interneurons^61^, had any effect on outward current amplitude (Fig. 2j, k). To confirm the specificity and potency of the interfering peptides used in hippocampal neurons we transfected plasmids coding for the last 11 C-terminal amino acids of either Goa, Gob and the scrambled version of Goa, into the Goa and Gob vectors in the TruPath assay used in Fig. 2h (Supplementary Fig. 7b). The interfering peptides reduced the activation of their cognate G protein in a dose-dependent manner, but had no effect on the alternate Go isoform. The scrambled peptide sequence had no effect on Goa or Gob activation.

Thus, adenosine-mediated membrane potential hyperpolarisation occurs mainly through A_1_R activation of Goa, in keeping with the high levels of expression of Goa *vs* Gob in the hippocampus^57^, and with the observation that the Goa-activating agonists adenosine, CPA and HOCPA (Fig. 2c-e, Supplementary Figs. 5 and 6) all induced membrane hyperpolarisation (Fig. 1h, i; Supplementary Figs. 2 and 5). Moreover, the absence of an effect of adenosine on membrane potential in Gz knockout mice^68^ argues against the possibility that the selective activation of Gz by adenosine observed in our PTX assays (Fig 2c, d; Supplementary Fig. 6) contributes to membrane hyperpolarisation. The data from recombinant receptors demonstrating the inability of BnOCPA to activate Goa (Fig. 2a, c-g) thus explains why BnOCPA did not cause membrane hyperpolarisation, and indeed prevented or reversed the hyperpolarisation induced by CPA or adenosine, respectively (Fig. 1h, i; Supplementary Fig. 2a, c).

### The Gα selectivity displayed by BnOCPA is reflected in non-canonical binding modes and a selective interaction with Gαi/o subunits

To better understand the unusual signalling properties of BnOCPA and the highly specific Gα coupling to Gob, we carried out dynamic docking simulations to study the basic orthosteric binding mode of BnOCPA in an explicit, fully flexible environment using the active cryo-EM structure of the A_1_R (PDB code 6D9H; Supplementary Movie 1). We previously reported that modifications at position N^6^ of the adenine scaffold modulated the agonist binding path to A_1_R^69^. More precisely, N^6^-cyclopentyl analogues (CPA and HOCPA) markedly interact with the extracellular loop 2 (ECL2) compared to adenosine, while BnOCPA (which bears the N^6^-cyclopentyl-2-benzyloxy group) is most prone to engage residues of the A_1_R located at the top of transmembrane helix 1 (TM1) and TM7. In the present study, we compared the bound-state BnOCPA to the non-Gα selective agonists adenosine and HOCPA, and an antagonist (PSB36) of the A_1_R (Fig. 3a-c). BnOCPA engaged the receptor with the same fingerprint as adenosine^70^ (Fig. 3a) and HOCPA (Fig. 3b, Supplementary Movie 2). Further explorations of the BnOCPA docked state using metadynamics (MetaD) simulations^71^ revealed interchangeable variations on this fingerprint (namely Modes A, B, and C; Fig. 3d-f; Supplementary Fig. 8) that could be distinguished by the orientation of the BnOCPA-unique benzyl group. Having established the possible BnOCPA binding modes, we examined the respective contribution of the orthosteric agonists, the G protein α subunit α5 (C-terminal) helix (GαCT), and the Gα protein subunit^72, 73^ to the empirically-observed G protein selectivity displayed by BnOCPA (Fig. 2a-g, Supplementary Fig. 6).

**Fig. 3.**
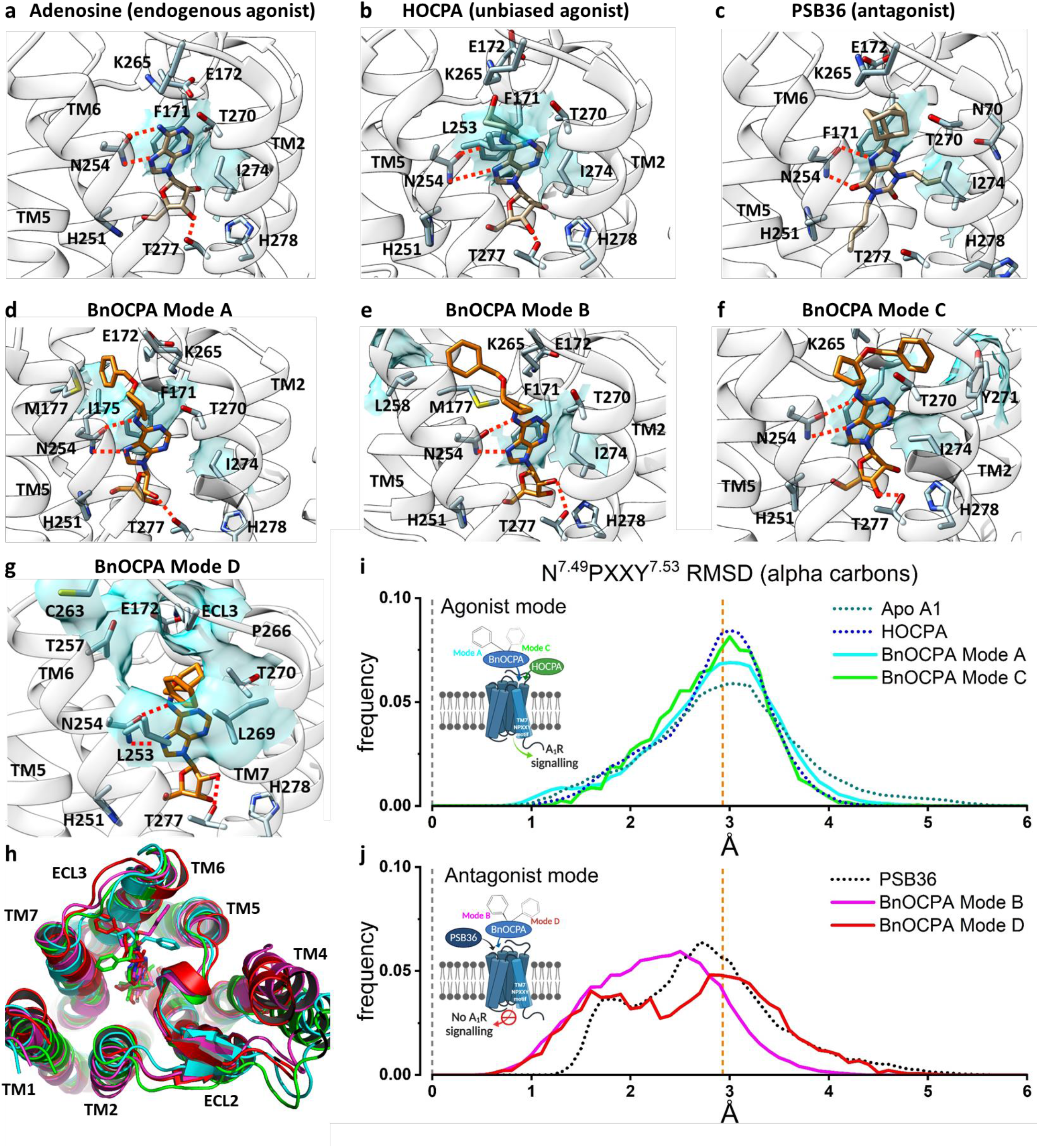
Molecular dynamics simulations reveal that BnOCPA binding modes can uniquely drive both agonist- and antagonist-like intracellular conformations of the A_1_R. a. Adenosine binding pose: N254^6.55^ (Ballesteros-Weinstein superscript enumeration) is engaged in key hydrogen bonds, while important hydrophobic contacts are shown as cyan transparent surfaces (F171^ECL2^ and I274^7.39^). **b** On the basis of structural similarities and the dynamic docking (Supplementary Movie 2), HOCPA was predicted to bind with a geometry analogous to adenosine; the cyclopentyl group makes further hydrophobic contacts with L253^6.54^, as shown by simulation. **c** The xanthine scaffold of the antagonist PSB36 makes hydrogen bonds with N254^6.55^ side chains and hydrophobic contacts with F171^ECL2^ and I274^7.39^. **d** BnOCPA agonist-like binding Mode A (Supplementary Movie 1): the benzyl group orients towards the ECL2 and makes hydrophobic contacts with I175^ECL2^ (and M177^5.35^) side chains. **e** BnOCPA antagonist-like binding Mode B: the benzyl group orients towards the top of TM5/TM6 and makes hydrophobic contacts with L258^6.59^ side chain. **f** BnOCPA agonist-like binding Mode C: the benzyl group orients towards the top of TM7 and makes hydrophobic contacts with Y271^7.36^ side chain. **g** Binding orientation of BnOCPA in antagonist Mode D: the benzyl group orients under ECL3 and occupies the hydrophobic pocket defined by L253^6.54^, T257^6.58^, T270^7.35^, and L269^7.34^. Key hydrogen bonds with N254^6.55^ and T277^7.42^ are shown as dotted lines; main hydrophobic contacts are highlighted as cyan transparent surfaces. **h** Extracellular view of the A_1_R showing the four BnOCPA binding Modes A (cyan), B (magenta), C (green), and D (red) as randomly extracted from the MD simulations. **i, j** Root-mean-square deviation (RMSD) distributions considering the inactive N^7.49^PXXY^7.53^ motif on the distal part of TM7 as reference. **i** HOCPA (blue broken line), BnOCPA Mode A (cyan curve), BnOCPA Mode C (green curve) and the apo receptor (dark green broken line) have a common distribution centring around the active confirmation of the A_1_R (orange broken line; Supplementary Fig. 9) leading to A_1_R signalling. In contrast, **j** PSB36 (black broken line), BnOCPA Mode B (magenta curve) and BnOCPA Mode D (red curve) RMSD values have the tendency to move closer to the inactive N^7.49^PXXY^7.53^ geometry (leftward shift of the curves towards broken grey line at x = 0) preventing A_1_R signalling.

#### Simulations in the absence of G protein

Firstly, following Dror et al.,^74^ we compared the dynamics of the BnOCPA-bound A_1_R with the corresponding dynamics of the receptor^75, 76^ bound to either HOCPA (Fig. 3b), the A_1_R antagonist PSB36 (Fig. 3c), or the apo receptor, our hypothesis being that there may be ligand-dependent differences in the way that the intracellular region of the receptor responds in the absence of the G protein. In these simulations the G protein was omitted so that inactivation was possible and so that the results were not G protein-dependent. The BnOCPA binding Modes A-C were interchangeable during MD simulations (Methods Table 1) but were associated with distinctly different dynamics, as monitored by changes in a structural hallmark of GPCR activation, the N^7.49^PXXY^7.53^ motif^77^ (Supplementary Fig. 9). Given the high flexibility shown by the BnOCPA benzyl group during the simulations and its lipophilic character, we hypothesized and simulated a further binding mode (namely Mode D) not explored during MD or MetaD simulations. This conformation involves a hydrophobic pocket underneath ECL3 (Fig. 3g) which is responsible for the A1/A2A selectivity^70^. Superimposition of the four BnOCPA binding Modes A-D reveals the highly motile nature of the benzyl group of BnOCPA (Fig. 3h) under the simulated conditions.

Quantification of the N^7.49^PXXY^7.53^ dynamics revealed that HOCPA, BnOCPA Mode A, BnOCPA Mode C and the apo receptor show a similar distribution of the RMSD of the conserved N^7.49^PXXY^7.53^ motif (Fig. 3i; Supplementary Fig. 9). In contrast, the non-canonical BnOCPA binding Modes B and D were responsible for a partial transition of the N^7.49^PXXY^7.53^ backbone from the active conformation to the inactive conformation (Supplementary Fig. 9) in a manner analogous with the antagonist PSB36 (Fig. 3j). Overall, the simulations revealed Mode D as the most stable BnOCPA pose (6.8 µs out 9 µs simulated starting from this configuration – Methods Table 1), while Mode B accounted for 3.6 µs out of 21 µs.

#### Dynamic Docking of GαCT

To simulate the agonist-driven interaction between the A_1_R and the G protein, the α5 (C-terminal) helix (GαCT) of the G protein (Gi2, Goa, Gob) was dynamically docked to the HOCPA- and BnOCPA-bound active A_1_R structure (again lacking G protein; Supplementary Movie 3). This allowed us to evaluate the effect of different GαCT on the formation of the complex with A_1_R to test the hypothesis that, of Goa, Gob and Gi2, only the GαCT of Gob would fully engage with the BnOCPA-bound active A_1_R, in line with the empirical observations of G protein selectivity summarized in Fig. 2c, d. Fig. 4a shows that the GαCT of Gob docked to the A_1_R via a metastable state (MS1) relative to the canonical state (CS1; Supplementary Movie 3), regardless of whether HOCPA or BnOCPA was bound. Fig. 4b, c show that the CS1 geometry corresponds to the canonical arrangement as found in the cryo-EM A_1_R:Gi protein complex, whereas state MS1 resembles the recently reported non-canonical state observed in the neurotensin receptor, believed to be an intermediate on the way to the canonical state^78^. In contrast, Fig. 4d-f show that the GαCT of Goa and Gi2 docks to the A_1_R to form metastable states MS2 and MS3. MS2 is similar to the β_2_-adrenergic receptor:GsCT fusion complex^79^, proposed to be an intermediate on the activation pathway and a structure relevant to G protein specificity. In this case however, it appears to be on an unproductive pathway.

**Fig. 4.**
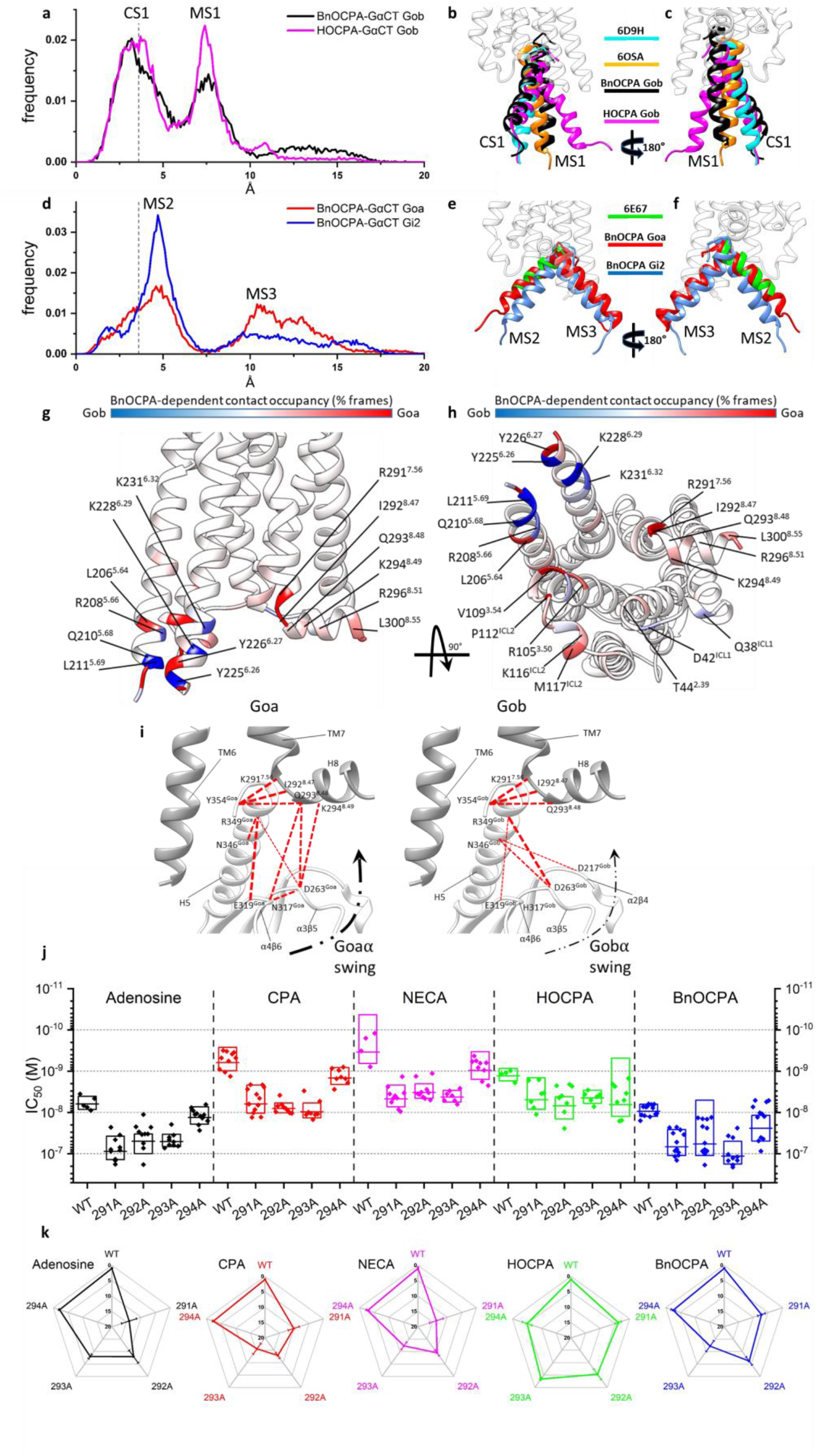
BnOCPA selectively induces canonical activation states at A_1_R:Gob, but non-productive metastable states at other Gαi/o subunits. a, b, c Dynamic docking of the Gob GαCT (last 27 residues) performed on the BnOCPA-A_1_R (black) and the HOCPA-A_1_R (magenta) complex, respectively. BnOCPA and HOCPA make productive couplings with the CT of Gob. **a** Frequency distribution of the RMSD of the last 15 residues of Gob GαCT (alpha carbon atoms) relative to the Gi2 GαCT conformation reported in the A_1_R cryo-EM structure 6D9H (the 3.6Å resolution of which is indicated by the dashed grey line): the two most probable RMSD ranges, namely canonical state CS1 and metastable state MS1, can be observed. **b, c** Two side views of representative MD frames of the most populated α5 clusters from the states CS1 and MS1. The last 15 residues of Gob GαCT in the CS1 states of both BnOCPA (black) and HOCPA (magenta) resemble the experimental Gi2 bound state (PDB code 6D9H – cyan). The alternative highly populated MS1 state is characterized by a binding geometry similar to the non canonical Gi intermediate state reported in the neurotensin receptor structure 6OSA (orange). **d, e, f** Dynamic docking of the Goa (red) and Gi2 (blue) GαCT (last 27 residues) performed on the BnOCPA-A_1_R complex. BnOCPA makes non-productive couplings with the CTs of Goa and Gi2. **d** Frequency distribution of the RMSD of the Goa (red) and Gi2 (blue) GαCT last 15 residues (alpha carbon atoms) relative to the Gi2 GαCT conformation reported in the A_1_R cryo-EM structure 6D9H (the resolution of which, 3.6Å, is indicated by the dashed grey line): the two most probable RMSD ranges are labelled as MS2 and MS3. **e, f** Two side views of representative MD frames of the most populated GαCT clusters from the states MS2 and MS3; the Goa (red) and Gi2 (blue) last 15 residues in the state MS2 overlap well with the putative Gs intermediate state (PDB code 6E67 – green). In the alternative highly populated state MS3, the GαCT helix orients in unique conformations that differ from those previously described. **g, h** For each residue the interaction plotted on the backbone is the difference between the Goa and Gob occupancies in the presence of orthosteric BnOCPA (% of MD frames in which interaction occurred). BnOCPA/A_1_R/Goa (inactive coupling) had the tendency to interact more with ICL2, TM3 TM7, and H8 (red), while BnOCPA/A_1_R/Gob (active coupling) formed more contacts with TM5 and TM6 (blue). **I** Residues in TM7 and H8 of the hA_1_R predicted by MD simulations to be of importance to A_1_R coupling to Goa (left) and Gob (right). **j, k**, Mutations of R291^7.56^, I292^8.47^, Q293^8.48^ and K294^8.49^ to alanine in the hA_1_R differentially affect agonist efficacy (**j**; IC_50_) against stimulated cAMP production. Data are shown for individual IC_50_ values from between 5 and 13 individual experiments, with the mean represented as the horizontal bar and ±1 SD represented as the box. The influence of the mutations can best be observed in the spider plot (**k**), which normalizes the reduction in IC_50_ for eachmutation and agonist relative to corresponding WT hA_1_R. The K294A mutation has little effect on agonist efficacy (< 5-fold change in IC_50_), while none of the mutations appreciably affect the efficacy of HOCPA. The R291A^7.56^, I292A^8.47^, and Q293A^8.48^ mutations strongly affect the efficacy of adenosine, CPA, NECA and BnOCPA.

#### MD simulations on the full G protein Gα subunit

To test the hypothesis that the non-functional BnOCPA:A_1_R:Goa complex showed anomalous dynamics, we increased the complexity of the simulations by considering the Gα subunit of the Goa and Gob protein bound to the A_1_R:BnOCPA (Mode B or D) complex or the Gob protein bound to A_1_R:HOCPA (a functional system). The most visible differences between Goa (Supplementary Movie 4) and Gob (Supplementary Movie 5) comprised the formation of transient hydrogen bonds between the α4-β6 and α3-β5 loops of Goa and helix 8 (H8) of the receptor (Supplementary Table 3). Similar contacts are present in the non-canonical state of the neurotensin receptor:Gi protein complex^78^. Overall, Goa interacted more with TM3 and ICL2 residues (Fig. 4g, h), while TM5 and TM6, along with ICL1, were more engaged by Gob (Fig. 4g, h). Interestingly, R291^7.56^ and I292^8.47^, which are located under the N^7.49^PXXY^7.53^ motif, showed a different propensity to interact with Goa or Gob. In this scenario, it is plausible that a particular A_1_R conformation stabilized by BnOCPA (as suggested by the simulations in the absence of G protein, Fig. 3i, j) may favor different intermediate states during the activation process of Goa and Gob.

To test the prediction from the MD simulations that R291^7.56^ and I292^8.47^ were involved in A_1_R/Gα coupling, we performed a series of site-directed mutagenesis (to alanine) against R291^7.56^, I292^8.47^ and the adjacent hydrophilic residues Q293^8.48^ and K294^8.49^ (Fig 4I) and compared the inhibition of forskolin-stimulated cAMP production in response to adenosine, CPA, NECA, HOCPA and BnOCPA in FlpIn-CHO cells against the wild-type (WT) hA_1_R (Fig 4J). Of these residues, none of which are reported to affect binding^80, 81^, K294^8.49^ had the least impact on potency. For the agonists, the mutations had minimal effects on HOCPA. In contrast A_1_R/Gα coupling induced by adenosine, CPA, NECA and BnOCPA was affected, but differentially so. These effects on potency (IC_50_ values) can be readily observed when individual mutant IC_50_ values are normalized to their respective WT controls (Fig. 4k), and revealed that R291^7.56^, I292^8.47^ and Q293^8.48^ are especially important for CPA and NECA coupling, R291^7.56^ for adenosine potency, and Q293^8.48^ for BnOCPA. These observations reinforce the MD simulations predictions related to H8 residues involved in G protein coupling of the agonist-stimulated A_1_R, and in particular suggest that R291^7.56^, I292^8.47^ and Q293^8.48^ are especially required for selective agonist coupling to Gαo/i, and may thus contribute to the Gα bias observed among these agonists (Fig. 2c, d). A more detailed analysis, involving saturation mutagenesis of these residues is required to provide a full characterization of their actions to direct agonist bias but is beyond the scope of this current study.

### BnOCPA does not depress heart rate, blood pressure or respiration: evidence for *in vivo* physiological selectivity at native A_1_Rs

Given BnOCPA’s clear differential effects in a native physiological system (Fig. 1), strong Gob selectivity (Fig. 2), unique binding characteristics (Fig. 3) and selective Gob interaction (Fig. 4), we hypothesised that these properties might circumvent a key obstacle to the development of A_1_R agonists for therapeutic use – their powerful effects in the cardiovascular system (CVS) where their activation markedly reduces both heart rate and blood pressure^12^. These cardiovascular effects are likely through Goa, which is expressed at high levels in the heart^82, 83^, particularly in the atria^84^, and which plays an important role in regulating cardiac function^85^. In contrast, and with parallels of differential Goa vs Gob expression in the hippocampus^57^, Gob may be absent or expressed at very low levels in the heart^84, 86^.Given this differential expression of Goa and Gob, and the lack of functional effect of BnOCPA on Goa (Fig. 2a-g), we predicted that BnOCPA would have minimal effects on the CVS. Moreover, given the antagonism of Goa-mediated actions by BnOCPA at native and recombinant A_1_Rs (Fig. 1h, i, Supplementary Fig. 2a, c, Fig. 2i), we further predicted that the actions of adenosine on the CVS may be attenuated by BnOCPA.

In initial experiments we screened BnOCPA for its effects on heart rate using an isolated frog heart preparation. In contrast to adenosine and CPA, which depress heart rate through hyperpolarisation caused by activation of cardiac sinoatrial K^+^ channels^87^, BnOCPA had no effect on heart rate, but markedly reduced the bradycardia evoked by adenosine (Supplementary Fig. 10a). Thus, BnOCPA appears not to activate A_1_Rs in the heart, but instead behaves like an antagonist in preventing the actions of the endogenous agonist. These observations have parallels with BnOCPA’s inability to activate A_1_Rs to hyperpolarise neurones, and indeed inhibiting or reversing the postsynaptic hyperpolarisation induced by typical A_1_R agonists (Fig. 1h, i; Supplementary Fig. 2a, c), and in preventing the A_1_R/Goa-mediated inhibition of cAMP accumulation by adenosine (Fig. 2i). Such antagonist-like behaviour may be explained by BnOCPA causing unique A_1_R conformations unlike those of conventional agonists (Fig. 3i, j), and driving non-canonical and ultimately non-productive interactions with Goa (Fig. 4).

To investigate the effects of BnOCPA in an intact mammalian system, we measured the influence of BnOCPA on heart rate and blood pressure in urethane-anaesthetised, spontaneously breathing adult rats. As expected, both resting heart rate and arterial blood pressure were significantly reduced by adenosine and CPA (Fig. 5a-d). In complete contrast, BnOCPA had no effect on either heart rate (Fig. 5a, c) or blood pressure (Fig. 5b, d), even when applied at two or three times the initial dose (Supplementary Fig. 11; Fig. 6e, f). These negative observations could not be explained by metabolism of BnOCPA to an inactive substance since BnOCPA is a very stable compound (half-life (t_1/2_) > 240 min in PBS at 37°C) with a human plasma stability of ∼100 % remaining after 120 mins suggesting a t_1/2_ > 240 min at 37°C. In addition, the *in vitro* metabolic t_1/2_ of BnOCPA was determined as > 60 mins using human liver microsomes (0.1 mg/mL, 37°C), and the intrinsic clearance (CL_int_) calculated as <115.5 μL/min/mg. This was in contrast to the reference compounds verapamil and terfenadine (0.1 μM) with t_1/2_ in human plasma determined as 33 and 10 min and CL_int_ as 213.1 and 683.0 μl/min/mg, respectively (see Supporting Data File 1). Further evidence that BnOCPA was present and active during these experiments was obtained from studies analogous to those in frog heart when BnOCPA was applied together with adenosine. In the intact anaesthetised rat, when co-applied with adenosine or CPA, BnOCPA abolished the bradycardia induced by both agonists, indicating its ability to bind to the A_1_R at the dose applied (Fig. 5a, c; Fig. 6g, Supplementary Figs. 10b and 11). Volumes of saline equivalent to the drug injections had no effect on either heart rate or blood pressure and there was no waning in the effects of adenosine responses with repeated doses (Supplementary Fig. 10c, d). Thus, BnOCPA does not appear to act as an agonist at CVS A_1_Rs, but instead antagonises the bradycardic effects of A_1_R activation on the heart.

Since adverse effects on respiration (dyspnea) limit the use of systemic A_1_R agonists^7^, we additionally examined the effects of BnOCPA on respiration. In urethane-anaesthetised, spontaneously breathing adult rats, intravenous injection of BnOCPA had no appreciable effect on respiration (Fig. 6a-d), even if the dose of BnOCPA was doubled or trebled (Fig. 6e, f). In stark contrast the selective A_1_R agonist CPA caused significant respiratory depression (Fig. 6a-d). Paralleling BnOCPA’s antagonism of adenosine- and CPA-induced depressions of heart rate (Fig. 5a, c; Supplementary Figs. 10b and 11), BnOCPA reduced the depression of respiratory frequency and minute ventilation caused by CPA (Fig. 6g, Supplementary Fig. 11). These data suggest that while BnOCPA targets and clearly engages the A_1_Rs responsible for adenosine and CPA’s cardiorespiratory depression, BnOCPA has no agonist action at these A_1_Rs.

**Fig. 5.**
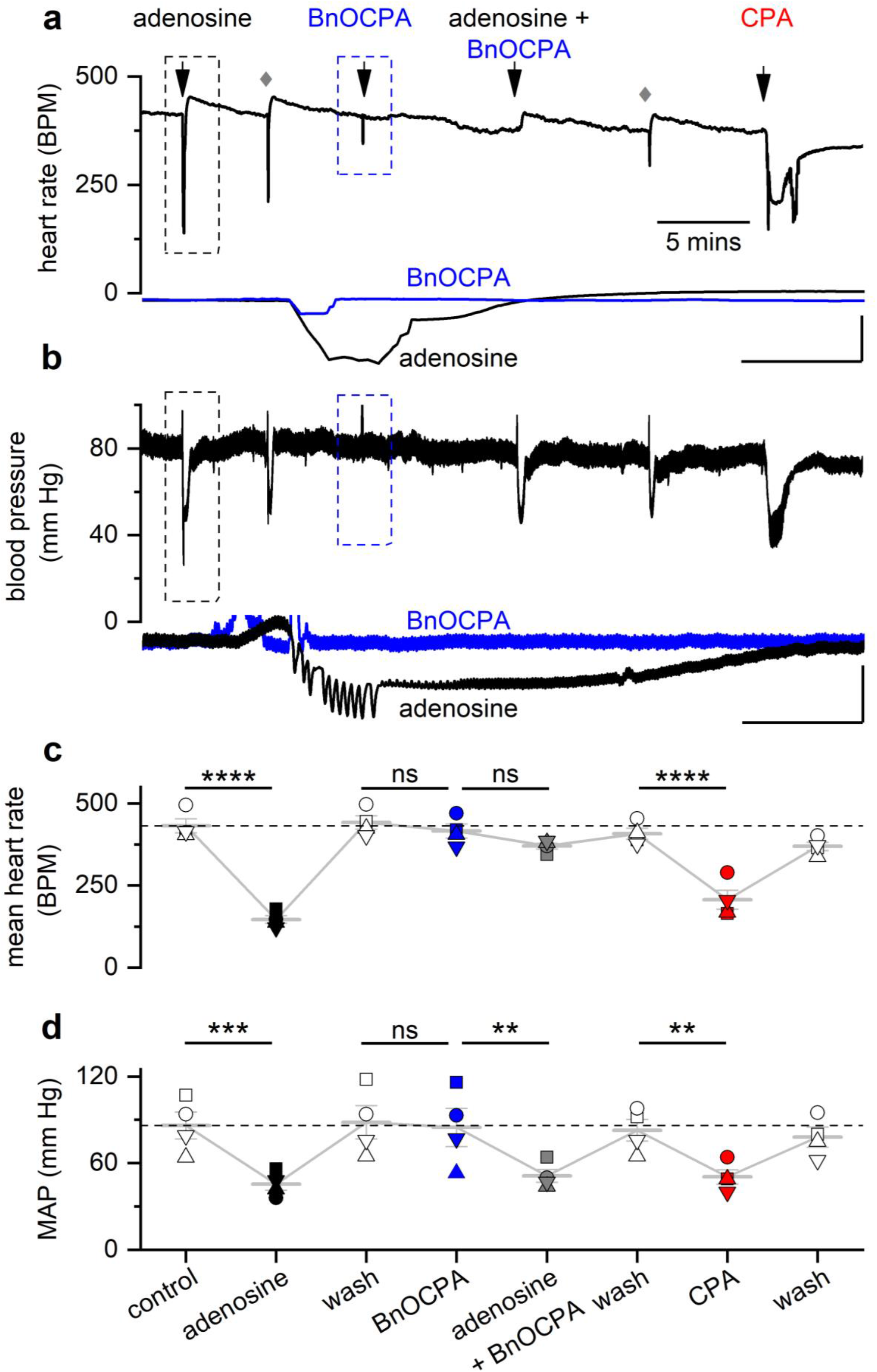
BnOCPA does not affect heart rate or blood pressure. a Examples of heart rate (HR) and **b** blood pressure traces from a single urethane-anaesthetised, spontaneously breathing rat showing the effects of adenosine (1 mg·kg^-1^), BnOCPA (8 µg·kg^-1^) and CPA (6 µg·kg^-1^). Adenosine, BnOCPA and CPA were all given as a 350 µL·kg^-1^ IV bolus. The intravenous cannula was flushed with 0.9% saline (grey diamonds) to remove compounds in the tubing. The overshoot in HR following adenosine applications is likely the result of the baroreflex. Insets are expanded HR and blood pressure responses to adenosine (black trace, boxed region in **a** and **b**) and BnOCPA (blue trace and boxed region in **a** and **b**). Scale bars measure: HR, 200 BPM and 6 s; blood pressure, 40 mm Hg and 6 s. **c, d** Summary data for 4 experiments. Data from each rat is shown as a different symbol. Means (± SEM, light grey bars) are connected to indicate the sequential nature of treatments across the four preparations. One-way RM ANOVA for: **c** HR, Greenhouse-Geisser corrected F(2.33, 7.00) = 68.27, P = 2.07 x10^-5^; **d** mean arterial blood pressure (MAP), Greenhouse-Geisser corrected F(1.84, 5.52) = 10.51, P = 0.014; with the following Bonferroni post hoc comparisons: The resting HR of 432 ± 21 BPM was significantly reduced to 147 ± 12 BPM (∼66 %, P = 2.76 x10^-11^) by adenosine. BnOCPA had no significant effect on HR (∼6%, 442 ± 20 vs 416 ± 21 BPM; P = 1) but prevented the bradycardic effects of adenosine (P = 2.71 x10^-9^ vs adenosine) when co-injected (mean change 51 ± 4 BPM; ∼12 %; P = 0.67). CPA significantly decreased HR (from 408 ± 17 to 207 ± 29 BPM; ∼50 %, P = 1.85 x10^-8^), a decrease that was not significantly different to the effect of adenosine (P = 0.12), but was significantly different to the effect of both BnOCPA (P = 9.00 x 10^-9^) and adenosine in the presence of BnOCPA (P = 6.69 x10^-7^). The resting MAP (86 ± 9 mm Hg) was significantly reduced by adenosine (∼47 %, 46 ± 4 mm Hg; P = 0.001). BnOCPA had no significant effect on its own on MAP (88 ± 11 vs 85 ± 13 mm Hg; P = 1) and did not prevent adenosine in lowering MAP to a value similar to adenosine on its own (51 ± 4 mm Hg; P = 1 vs adenosine; P = 0.012 vs BnOCPA alone). CPA significantly decreased MAP (from 83 ± 8 to 51 ± 5 mm Hg; P = 0.017), a decrease that was not significantly different to the effect of adenosine in the absence or presence of BnOCPA (P = 1 for both). ns, not significant; **, P < 0.02; ***, P < 0.001; ****, P < 0.0001.

**Fig. 6.**
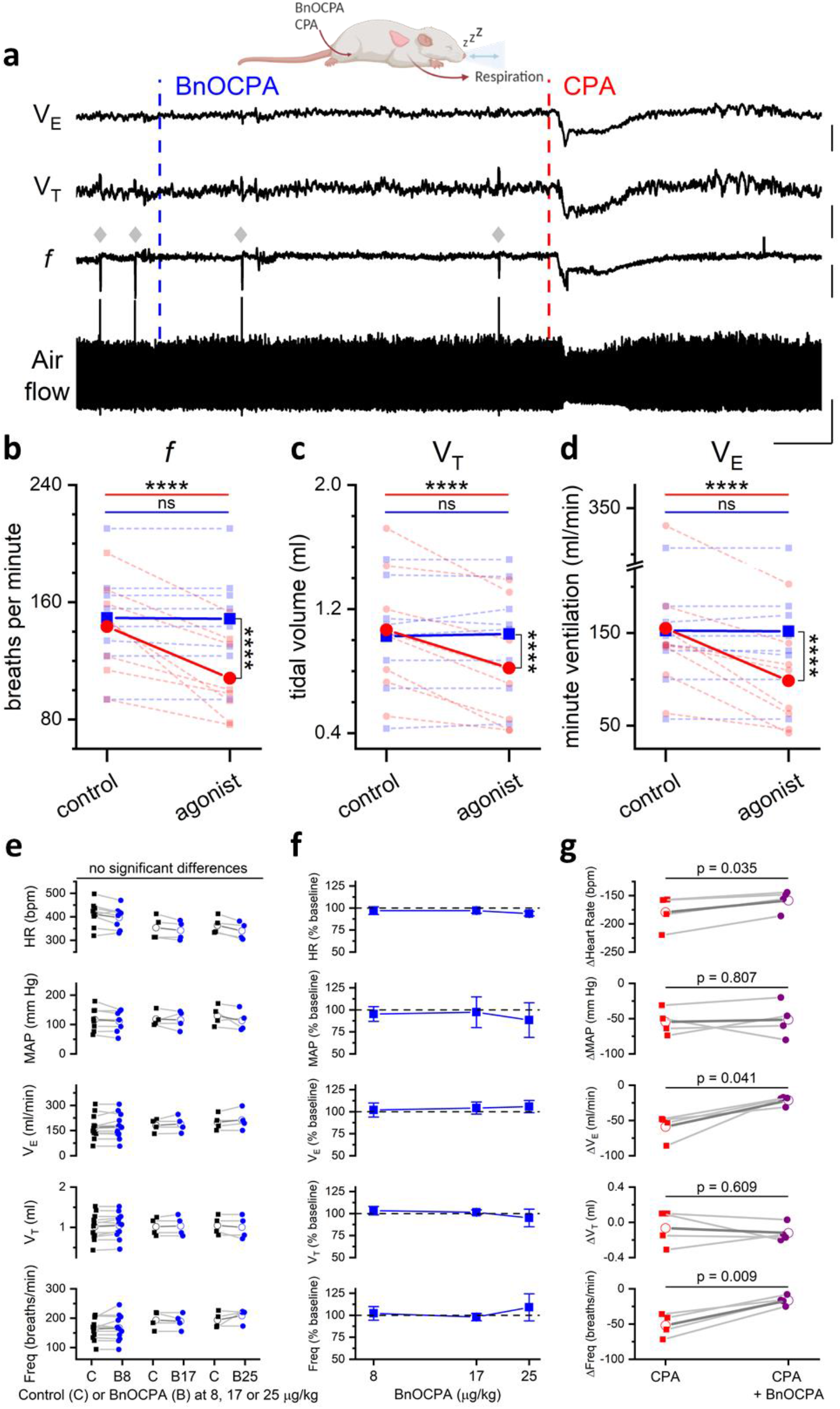
BnOCPA does not cause respiratory depression. **a** examples of tracheal airflow, respiratory frequency (*f*), tidal volume (V_T_) and minute ventilation (VE) from a single urethane-anaesthetised, spontaneously breathing rat showing the lack of effect of BnOCPA on respiration and the respiratory depression caused by CPA. BnOCPA and CPA were given as a 350 µL·kg^-1^ IV bolus at the times indicated by the vertical broken lines (BnOCPA, 8 μg/kg, blue; CPA, 6 µg·kg^-1^, red). Grey diamonds indicate spontaneous sighs. Scale bars measure: 180 s and: airflow, 0.5 mL; *f*, 50 breaths per minute (BrPM); V_T_, 0.25 mL; V_E_, 50 mL/min. **b, c, d** Summary data for 8 anaesthetised rats. Data from each rat is shown before and after the injection of BnOCPA (blue squares and broken lines) and CPA (red circles and broken lines) together with the mean value for all animals (solid lines) for *f*, V_T_ and V_E_, respectively. One-way RM ANOVA: For: **b**, *f*, Greenhouse-Geisser corrected F(1.20, 8.38) = 30.4, P = 3.48 x 10^-4^; **c**, V_T_, F(3, 21) = 15.9, P = 1.25 x 10^-5^, and **d**, VE, Greenhouse-Geisser corrected F(1.19, 8.34) = 15.77, P = 0.003, with the following Bonferroni post hoc comparisons: Following BnOCPA, *f* (149 ± 12 BrPM), V_T_ (1.0 ± 0.1 mL), and V_E_ (152 ± 26 ml/min) were not altered (P = 1) compared to resting values *f* (149 ± 12 BPM), V_T_ (1.0 ± 0.1 mL), and V_E_ (152 ± 26). In contrast to CPA, which reduced *f* (108 ± 10 BrPM), V_T_ (0.8 ± 0.1 mL), and V_E_ (99 ± 19 ml/min) compared to resting values *f* (143 ± 11 BrPM; p = 4.05 x 10^-6^), V_T_ (1.1 ± 0.1 mL; P = 2.58 x10^-5^), and V_E_ (155 ± 28; P = 5.52 x 10^-5^). Whilst the control resting values before administration of BnOCPA and CPA were not different to one another (P = 1). The effects of CPA were significantly greater than BnOCPA for *f* (P = 4.48 x 10^-7^), V_T_ (P = 1.15 x10^-4^), and V_E_ (P = 1.16 x10^-4^). Horizontal significance indicators above the data show differences between resting values and following IV administration of either BnOCPA (blue line) or CPA (red line). Vertical significance indicators show differences between the effects of BnOCPA and CPA. **e,** Individual data for the three doses of BnOCPA (blue circles) compared to their preceding baseline (black squares). The mean is shown as an open symbol. One-way ANOVA with Bonferroni corrections found no differences in: HR (p = 0.07), MAP (p = 1), Freq (p = 0.2), V_T_ (p = 1), or V_E_ (p = 0.9). **f**, Average data from the four animals in **e** showing cardiorespiratory variables as a percentage of their preceding baseline and as a function of increasing dose of BnOCPA (log10 scale). **g**, Individual data from four rats showing the effect (difference from previous baseline) of CPA in the absence (red squares) and presence (purple circles) of BnOCPA (8 µg/kg). The mean is shown as an open symbol. Paired t-tests indicated a significant reduction in the effects of CPA by BnOCPA on HR (CPA: 179 ± 15 bpm vs BnOCPA: 159 ± 10 bpm; p = 0.035), V_E_ (CPA: 59 ± 9 ml/min vs BnOCPA: 21 ± 3 ml/min; p = 0.041) and Freq (CPA: 52 ± 8 breaths/min vs BnOCPA: 17 ± 3 breaths/min; p = 0.009), with no change in: MAP (p = 0.807) or V_T_ (p = 0.609). Data is shown as mean ± SEM. Raw traces from a representative experiment can be found in Supplementary Fig. 11

### BnOCPA is a potent analgesic

Our observations of a lack of effect of BnOCPA on the CVS and respiration prompted an investigation into a potential application of A_1_R agonists that had previously been severely curtailed by adverse cardiorespiratory events^7, 16^, namely the use of A_1_R agonists as analgesics. Since sedation or motor impairment can be mistaken for analgesia, we tested BnOCPA in a sensitive assay for balance and motor coordination, the rotarod, in which the ability of a rodent to remain upon a slowly accelerating rotating cylinder is a measure of alertness and motor function. As a positive control for the sensitivity of the test, we showed that the ability of animals treated with morphine to remain on the rotating cylinder was strongly impaired (Fig. 7a). In contrast, the performance of animals treated with BnOCPA, delivered either intravenously or intraperitoneally, was indistinguishable from vehicle-treated mice (Fig. 7a). This was true even if BnOCPA was injected intravenously at three times the dose (Fig. 7a), which, while having no cardiorespiratory actions on its own, prevented cardiorespiratory depression caused by adenosine and CPA (Figs. 5 and 6; Supplementary Figs. 10 and 11). Thus, BnOCPA does not induce sedation or locomotor impairment that could confound interpretations of analgesia.

**Fig. 7.**
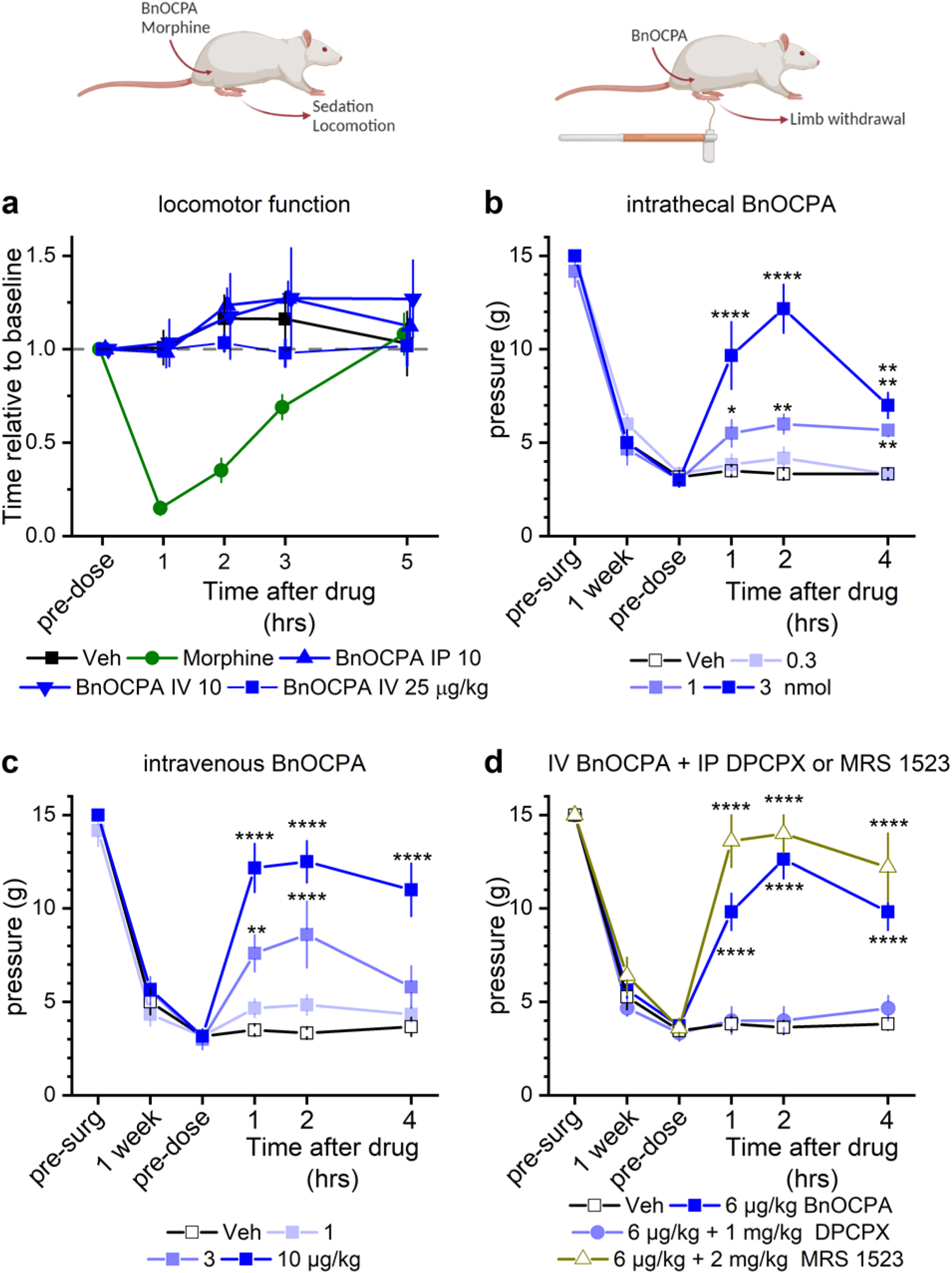
BnOCPA is a potent analgesic without causing sedation. a. BnOCPA does not induce sedation or affect motor function. BnOCPA was administered intravenous (IV; n = 6) or intraperitoneally (IP; n = 6) at 10 µg/kg as per the maximum dose used in the neuropathic pain study (Fig. 7b), and at 25 µg/kg IV (n = 6), the highest dose used in the cardiorespiratory experiments (Fig. 6; Supplementary Fig 11). Morphine (n = 6) was administered at 15 mg/kg subcutaneously as a positive control for sedation and motor impairment. Saline (Veh; n = 6) was administered subcutaneously at the same volume as the morphine injection. Rats were tested on the rotarod over a period of 5 hours after injection. BnOCPA did not affect motor function at analgesic or higher doses. Data points are normalised to pre-dose performance to take into account individual differences and are offset for clarity. **b, c** BnOCPA alleviates mechanical allodynia in a spinal nerve ligation (Chung) model of neuropathic pain when administered **b** via an intrathecal (IT) or **c** an IV route. Prior to surgery (pre-surg) animals had similar sensitivity to tactile stimulation as assessed by Von Frey hair stimulation. Spinal nerve ligation subsequently caused hypersensitivity to touch (mechanical allodynia) as evidenced by the reduction in the tactile pressure necessary to elicit paw withdrawal (paw withdrawal threshold; PWT) at 1 week after surgery. PWT reaches a similar nadir across all groups prior to vehicle or BnOCPA infusion (pre-dose). Administration of BnOCPA significantly increased PWT in the limb ipsilateral to the site of injury, in a dose-dependent manner (one-way ANOVA (pre-dose, 1, 2 and 4 hrs) for IT BnOCPA F(3,88) = 21.9, P = 1.10 x 10^-10^; for IV BnOCPA F(3,92) = 18.1, P = 2.70 x 10^-9^). Fisher LSD post-hoc comparisons showed significant differences at: IT 1 nmol at 1 and 2 hrs, P = 0.001 and 4.16 x 10^-5^, respectively, and 3 nmol at 1, 2 and 4 hrs, P = 9.52 x 10^-11^, 1.42 x 10^-11^ and 1.41 x 10^-8^, respectively; IV 3 µg/kg at 1, 2 and 4 hrs, P = 0.044, 0.008 and 0.019, respectively, and 10 µg/kg at 1, 2 and 4 hrs, P = 1.37 x 10^-8^, 6.81 x 10^-14^ and 3.23 x 10^-4^, respectively. **b** and **c**, n = 6 per treatment, except for 1 nmol BnOCPA, n = 5. **d** The analgesic effects of BnOCPA (6 µg/kg IV) were prevented by the A_1_R antagonist DPCPX (1 mg/kg IP), but not the A_3_R-selective antagonist MRS 1523 (2 mg/kg IP). Post-hoc LSD comparisons across all four groups and four time points (pre-dose, 1, 2 and 4 hrs; F(15,116) = 26.8, P = 0) revealed that BnOCPA at 6 µg/kg (IV) elicited significant analgesia compared to vehicle-treated animals at 1, 2, and 4 hours post-dosing (P = 4.69 x 10^-9^, 3.50 x 10^-16^, 4.69 x 10^-9^, respectively), which persisted in the presence of the selective A_3_R antagonist MRS 1523 over the same time period (P = 4.42 x 10^-13^, 3.38 x 10^-14^, 1.81 x 10^-10^, respectively). In contrast, the PWT in DPCPX-treated animals did not differ from those in the vehicle group (P = 0.872, 0.748, 0.453 at 1, 2 and 4 hours, respectively). n = 11 for BnOCPA and vehicle groups; n = 6 for the DPCPX group and n = 5 for the MRS 1523 group. Averaged data is presented as mean ± SEM. ns, not significant; *, P < 0.05; **, P < 0.02; ***, P < 0.001; ****, P < 0.0001.

To assess the potential of BnOCPA as an analgesic, we used a rat model of chronic neuropathic pain (spinal nerve ligation)^88^ a feature of which is mechanical allodynia whereby the affected limb is rendered sensitive to previously innocuous tactile stimuli, and which models the debilitating human clinical condition of chronic pain, which affects between 20 and 50% of the population^89, 90^, and which carries a major global burden of disability^91^.

Both intrathecal (Fig. 7b) and intravenous (Fig. 7c) BnOCPA potently reversed mechanical allodynia in a dose-dependent manner. Thus, BnOCPA exhibits powerful analgesic properties at doses devoid of sedative or cardiorespiratory effects, and at several orders of magnitude lower than the non-opioid analgesics pregabalin and gabapentin^92^. To test if this analgesia was mediated by the activation A_1_Rs by BnOCPA, we used the selective A_1_R antagonist, DPCPX. Prior administration of DPCPX prevented the reversal of mechanical allodynia by BnOCPA (Fig. 7d), confirming the importance of A_1_Rs in mediating the analgesic actions of BnOCPA. In contrast, the rat A_3_R-selective antagonist MRS 1523^93, 94^, which is effective in reversing analgesia caused by selective A_3_R agonists^95–97^, had no effect on the analgesia caused by BnOCPA, and indeed may have provoked a slight facilitation. This may be due to the reported elevated adenosine tone^98^ and activation of A_3_Rs^99^ in the neuropathic spinal cord, which may have resulted in the desensitization of A_1_R-mediated responses^100^. These observations confirm that the analgesia provoked by BnOCPA is mediated via the selective activation of A_1_Rs.

## Discussion

Biased agonists at GPCRs offer great potential for the preferential activation of desirable intracellular signalling pathways, while avoiding, or indeed blocking those pathways that lead to adverse or unwanted effects^3, 27^. While this, and the potential to exploit previously unattractive drug targets such as the A_1_R, have been appreciated, translation of *in vitro* observations, particularly of Gα bias, to native receptors *in vivo* has been problematic^3, 4, 27^. Here we have shown that translation of *in vitro* selectivity among Gα subunits, identified using two separate assays, to an intact physiological system is possible through a benzyloxy derivative (BnOCPA) of the selective A_1_R agonist CPA. Moreover, this Gα selectivity has occurred in the context of the A_1_R, an attractive, but notoriously intractable drug target by virtue of the profound cardiorespiratory consequences of its activation by conventional A_1_R agonists.

BnOCPA was first reported as a final compound in a patent, where it was described to be selective for the A_1_R with respect to its binding affinity, and in reducing elevated intraocular pressure in the treatment of glaucoma or ocular hypertension^34^. We have previously prepared BnOCPA (and HOCPA)^33^ for assessment as part of our synthetic campaign to develop potent and A_1_R-selective fluorescent ligands. The N^6^-substituent (1*R*,2*R*)-2-aminocyclopentan-1-ol) present in BnOCPA and HOCPA is also found in the experimental and later discontinued^101^ drug CVT-3619 (later named GS 9667), which has been described as a partial, selective agonist of the A_1_R and shown to reduce cAMP content and consequently lipolysis in rat adipocytes^102^.

Having identified BnOCPA as a selective Gob agonist at recombinant A_1_Rs *in vitro*, we established that this unusual property can be translated into the selective activation of native A_1_Rs in both the *in vitro* CNS and *in vivo* cardiorespiratory and peripheral nervous systems. Moreover, these properties of BnOCPA were observed at A_1_Rs expressed by three different species: amphibian, rat, and human. While BnOCPA bound to and induced A_1_R coupling to Gαi/o subunits recruited by prototypical A_1_R agonists such as adenosine and CPA, BnOCPA selectively activated Gob among the six Gαi/o subunits. This likely reflected BnOCPA’s non-canonical binding profile at the A_1_R, which had profound implications for the interaction with the GαCT in terms of different binding pathways and intermediate states, and in the different intra- and intermolecular hydrogen bond patterns and contacts observed in the simulations of the A_1_R in complex with either Goa (inactive) or Gob (active). Predictions from the MD simulations suggested four hitherto uncharacterised residues as being important for the interaction between the A_1_R and Gαi/o. Individual mutations in three of these contacts, R291^7.56^, I292^8.47^, Q293^8.48^, differentially impacted agonist efficacy, with adenosine and HOCPA being relatively unaffected compared to the stronger effects on the efficacy of CPA, NECA and BnOCPA. These and other molecular differences in the coupling of the A_1_R to Gαi/o are likely to underlie the ability of the BnOCPA-bound A_1_R to selectively trigger Gob activation among the six Gαi/o subtypes.

The unique and unprecedented Gα selectivity displayed by BnOPCA has physiological importance since it is able to inhibit excitatory synaptic transmission without causing neuronal membrane hyperpolarisation, sedation, bradycardia, hypotension or dyspnea. BnOCPA thus overcomes cardiovascular and respiratory obstacles to the development of adenosine-based therapeutics that have plagued the field since their first description nine decades ago^103^. As a first, but significant, step towards this, we demonstrate that BnOCPA has powerful analgesic properties via A_1_Rs in an *in vivo* model of chronic neuropathic pain, potentially through a mechanism which may involve a combination of inhibition of synaptic transmission in peripheral and spinal pain pathways, and the hyperpolarisation of Gob-containing nociceptive neurons. Chronic pain, a condition that a large proportion of the population suffers on a constant or frequent basis^89, 90^ and associated with a major global burden of disability^91^ is, however, a disorder for which the current treatments are either severely lacking in efficacy^104^ or, in the case of opioids, come with unacceptable harms such as adverse gastrointestinal effects, respiratory depression, tolerance, dependence and abuse potential^105^. Accordingly, novel treatments for chronic pain are urgently required.

We have shown that highly selective Gα agonism *in vitro* can be translated into selective activation of native A_1_Rs to mediate differential physiological effects, and have identified a novel molecule capable of doing so. We have also explored molecular mechanisms by which this could occur, and demonstrated pain as one potential and wide-reaching therapeutic application. Such discoveries are of importance in both understanding GPCR-mediated signalling, and in the generation of both new research tools and therapeutics based on the untapped potential of biased, and indeed Gα-selective, agonists such as BnOCPA.

## Acknowledgements

We gratefully acknowledge the support of the University of Warwick (URSS Awards to SH and JO; Warwick Ventures Proof of Concept Fund awards to MJW & BGF), the Leverhulme Trust (RPG-2017-255, CAR and GL to fund KB and GD), the BBSRC (BB/M00015X/2, GL and BB/M01116X/1, PhD studentship to EH), the MRC (MR/J003964/1; IW) and The Swiss National Science Foundation (PP00P2_123536 and PP00P2_146321, MLo). AS is supported by a European Scholarship from the Cambridge Trust, SC is funded by an AstraZeneca PhD studentship and XH is funded by a China Scholarship Council Cambridge International Scholarship. RH is supported by an MRC Discovery Award (MC_PC_15070). CAR is a Royal Society Industry Fellow. We would like to thank: Stephen Hill, Stephen Briddon and Mark Soave (University of Nottingham) for gifting the Nluc-tagged adenosine receptor cell lines, the fluorescent antagonist AV039 and technical advice; Kathleen Caron and Duncan Mackie (University of North Carolina) for the β-arrestin1/2-YFP constructs, and Annette Gilchrist (Midwestern University) and Heidi Hamm (Vanderbilt University) for assistance with the Gαo interfering peptide plasmids. We are grateful to: Kevin Moffat and the Biochemistry students of the School of Life Sciences at the University of Warwick for access to their frog heart preparations; Nick Dale, Mark Wigglesworth, Jens Kleinjung for discussions and comments on draft manuscripts, and Arthur Christopoulos for a pre- publication copy of the adenosine A1 cryo-EM structure. *In vivo* studies on neuropathic pain were funded and undertaken by NeuroSolutions Ltd. Illustrative figures were made in ©BioRender – biorender.com. Venn diagram made at http://bioinformatics.psb.ugent.be/webtools/Venn/.

## Author Contributions

Experiments were designed by MJW, RH, GD, CAR, GL, FYZ, WI, DSp, BGF, and were performed by MJW, EH, RH, KB, IW, SC, AS, HFW, DSa, XH, WI, CLM, ED, CH, SH, JO.

Compounds were synthesised by MLe, BP, Mlo. Molecular dynamics and docking simulations were designed and carried out by GD and CAR. Work was originally conceived by MJW and BGF. The manuscript was written by MJW, GD, CAR, GL, BGF, with valuable input from EH, RH, KB, Mlo, IW, WI and DSp.

## Data Availability

The data and materials that support the findings of this study are available from the corresponding author upon reasonable request.

## Supplementary Data

### Supplementary Figures

**Supplementary Fig. 1.**
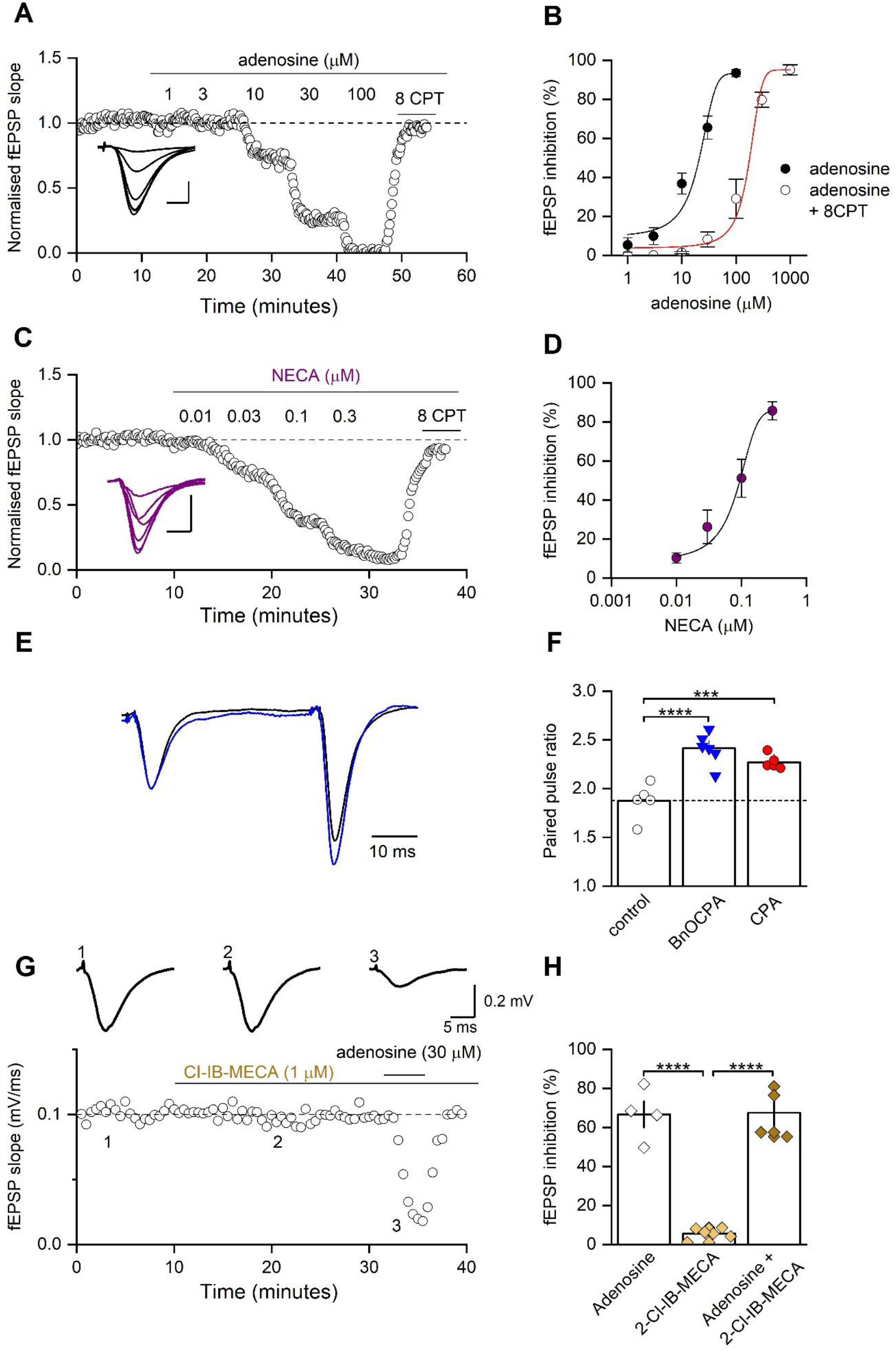
A_1_R, but not A_3_R, agonists inhibit excitatory synaptic transmission at hippocampal synapses. **A**, Increasing concentrations of adenosine reduced fEPSP slope, an effect reversed by the A_1_R antagonist 8CPT (2 µM). Inset, superimposed fEPSP averages in control and in increasing concentrations of adenosine. Scale bar measures 5 ms and 0.25 mV. **B**, Concentration-response curve for adenosine (IC_50_ = 20 ± 4.3 µM, *n* =11 slices) and for adenosine with 2 µM 8CPT (IC_50_ = 125 ± 10 µM *n* = 5 slices). **C**, Increasing concentrations of the A_1_R agonist NECA reduced fEPSP slope, an effect reversed by 8CPT (2 µM). Inset, superimposed fEPSP averages in control and in increasing concentrations of NECA. Scale bar measures 5 ms and 0.25 mV. **D**, Concentration-response curve for NECA (IC_50_ = 8.3 ± 3 nM, *n* =11 slices). **E**, Example of average (5 traces) superimposed paired-pulse fEPSP waveforms (50 ms inter-pulse interval) in control (black trace) and in the presence of BnOCPA (100 nM; blue trace). The fEPSP waveforms have been normalised to the amplitude of the first fEPSP in control. BnOCPA increased paired-pulse facilitation, indicative of a BnOCPA-induced reduction in the probability of glutamate release. **F**, Data summary. For a paired-pulse interval of 50 ms, the paired-pulse ratio was significantly increased (one-way ANOVA; F(2, 14) = 21.72; P = 5.11 x 10^-5^) from 1.88 ± 0.07 in control (*n* = 6 slices) to 2.41 ± 0.07 in BnOCPA (*n* = 6 slices, P = 5.17 x 10^-5^) and 2.27 ± 0.03 in CPA (60 nM; *n* = 5, P = 0.001). **G**, The potent and selective A_3_R agonist 2-Cl-IB-MECA had no effect on the fEPSP even at a high concentration (1 µM) and did not prevent adenosine (30 µM) from inhibiting synaptic transmission to an extent comparable to that seen in the absence of 2-Cl-IB-MECA (Panels A, B). Data presented shows the time course of an exemplar experiment with the inset fEPSPs taken at the times indicated (1) before, (2) during 2-Cl-IB-MECA, and (3) during adenosine application in the continued presence of the selective A_3_R agonist. **H**, Summary for the effects of 2-Cl-IB-MECA (1 µM) on fEPSPs and on the depression caused by adenosine (*n* = 4 – 6 slices; one-way ANOVA; F(2, 11) = 65.60; P = 7.71 x 10^-7^). Adenosine (30 µM) and adenosine (30 µM) in the presence of 2-Cl-IB-MECA (1 µM) depressed the fEPSP to comparable levels (66.7 ± 6.7 % and 67.5 ± 6.5 %; P = 1), and to a significantly greater extent than that caused by 2-Cl-IB-MECA (5.6 ±1.1 %; P = 3.63 x 10^-6^ vs adenosine, and P = 3.17 x 10^-6^ vs adenosine plus 2-Cl-IB MECA. Averaged data is presented as mean ± SEM. ***, P < 0.001; ****, P < 0.0001.

**Supplementary Fig. 2.**
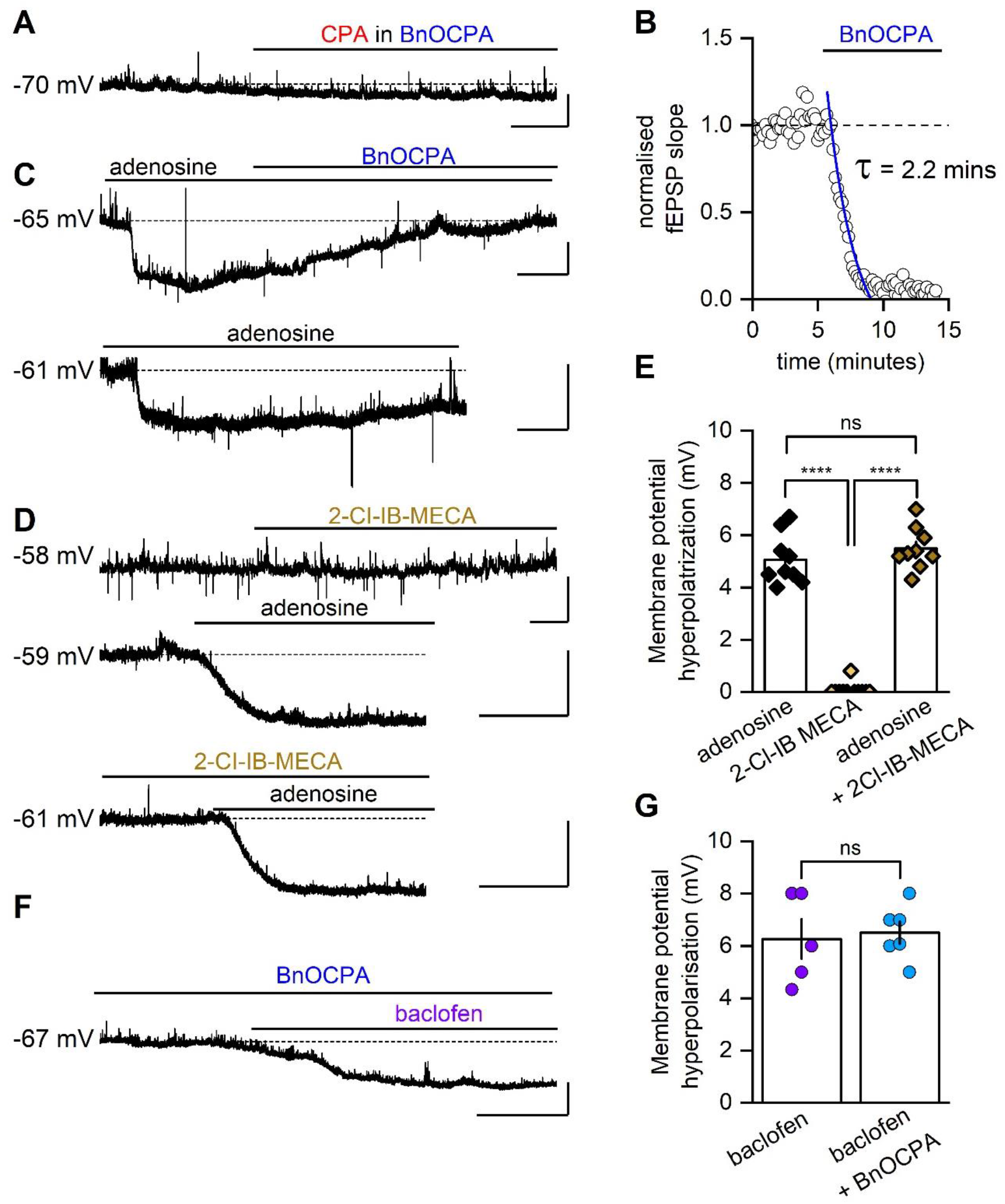
BnOCPA, but not the A_3_R agonist 2-Cl-IB-MECA, selectively inhibits membrane hyperpolarisation induced by prototypical A_1_R agonists. **A**, Membrane potential trace recorded from a CA1 pyramidal cell. BnOCPA (300 nM) reduced the effect of CPA (300 nM; quantified in main text Fig. 1i). **B,** The same solution of BnOCPA (300 nM), which had no effect on membrane potential, abolished synaptic transmission in a sister slice (inhibition fitted with a single exponential; τ = 2.2 mins). **C**, BnOCPA reversed the hyperpolarising effect of adenosine (100 µM; similar observations were made in 3 other cells), which (lower trace) cannot be accounted for by fatigue of adenosine-mediated hyperpolarisation (similar observations of sustained hyperpolarisations to adenosine were made in 2 other cells). **D**, 2-Cl-IB-MECA had no effect on membrane potential even when applied at a high concentration (1 µM). Moreover, the membrane hyperpolarisation caused by adenosine (30 µM) was not affected by prior incubation and co-application with 2-Cl-IB-MECA (1 µM). **E**, Summary of data from 9 – 11 cells showing that 2-Cl-IB-MECA does not affect membrane potential, nor does it prevent adenosine from inducing membrane hyperpolarisation, in contrast to BnOCPA, which reverses adenosine-mediated membrane hyperpolarisation. Bonferroni post-hoc comparisons after a one-way ANOVA (F(2,26) = 183.83, P = 4.441 x 10^-16^) showed no significant (ns) difference between adenosine application in the absence or presence of 2-Cl-IB-MECA (P = 0.621), but significantly smaller hyperpolarisations to 2-Cl-IB-MECA compared to adenosine alone (P = 3.029 x 10^-14^), or adenosine in the presence of 2-Cl-IB-MECA (P = 4.177 x 10^-15^). ****; P < 0.0001. **F,** Application of baclofen (10 µM) in the presence of BnOCPA (300 nM) hyperpolarised the membrane potential (from -67 to -74 mV). Scale bars measure 5 mV and 25 s (2-Cl-IB-MECA) 50 s (CPA), 200 s (adenosine) or 100 s (baclofen). **G**, Data summary of baclofen/BnOCPA experiments. The mean hyperpolarisation produced by baclofen in the presence of BnOCPA was not significantly different (ns; unpaired t-test) from that produced by baclofen in control conditions (6.5 ± 0.43 mV vs 6.3 ± 0.76 mV, P = 0.774, *n* = 5 – 6 cells for each condition). Bar chart displays individual data points and mean ± SEM.

**Supplementary Fig. 3.**
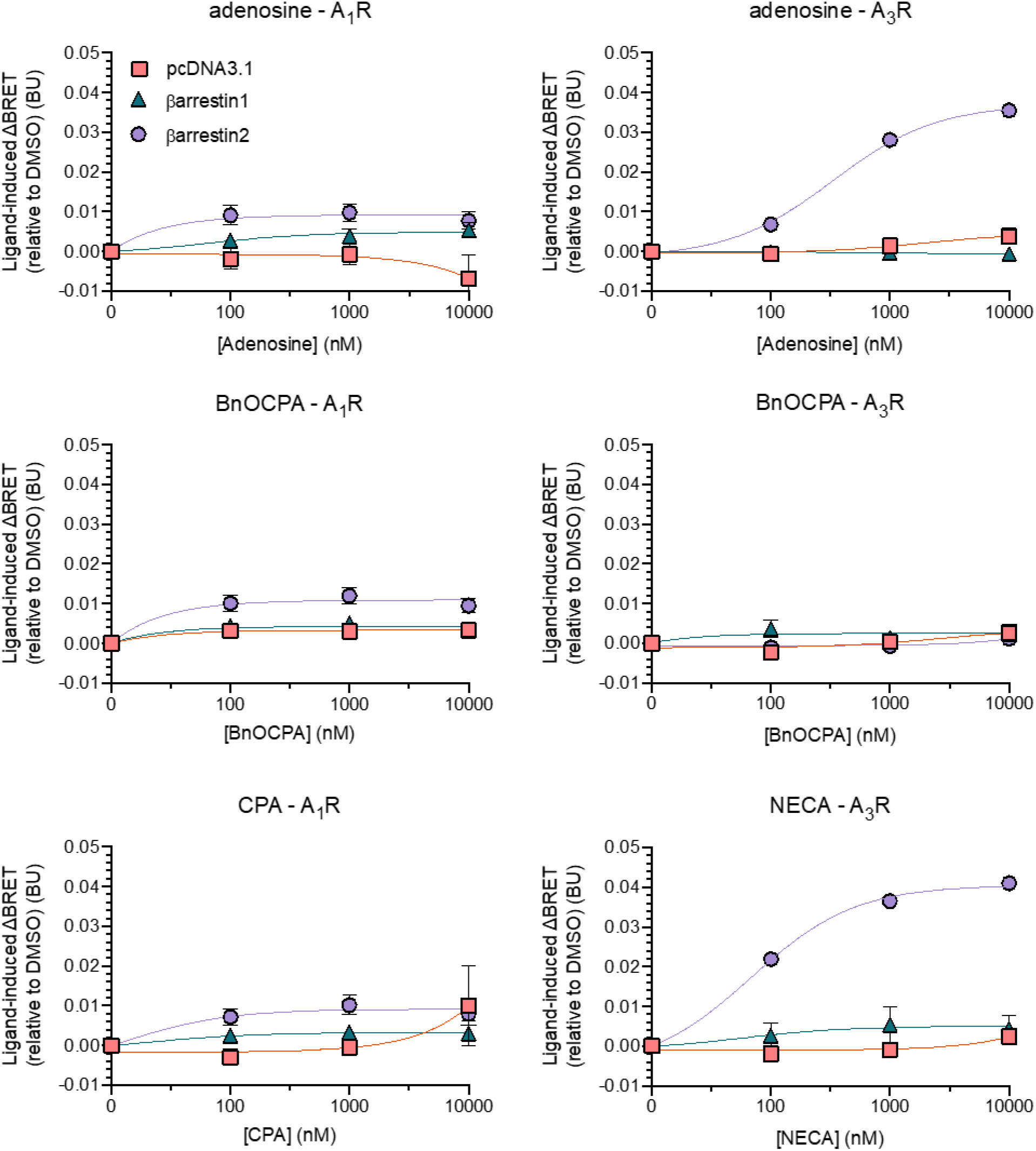
β-arrestin1 or β-arrestin2 recruitment to the human A_1_R or A_3_R. Interactions were detected via BRET using a C-terminally Nluc-tagged GPCR (A_1_R, left panels, or A_3_R, right panels) and C-terminally YFP-tagged β-arrestin1 or β-arrestin2, or pcDNA3.1 (negative control). Cells were stimulated with 3 agonists (top panels – adenosine; middle panels – BnOCPA; lower left – CPA; lower right – NECA). Note lack of either β-arrestin1 or β-arrestin2 recruitment to the A_1_R either by adenosine, CPA or BnOCPA, which yield BRET signals comparable to the vector control experiments (pcDNA3.1; top panels). A_3_R recruitment of β-arrestin2 is provided as a positive control for the BRET assay.

**Supplementary Fig. 4.**
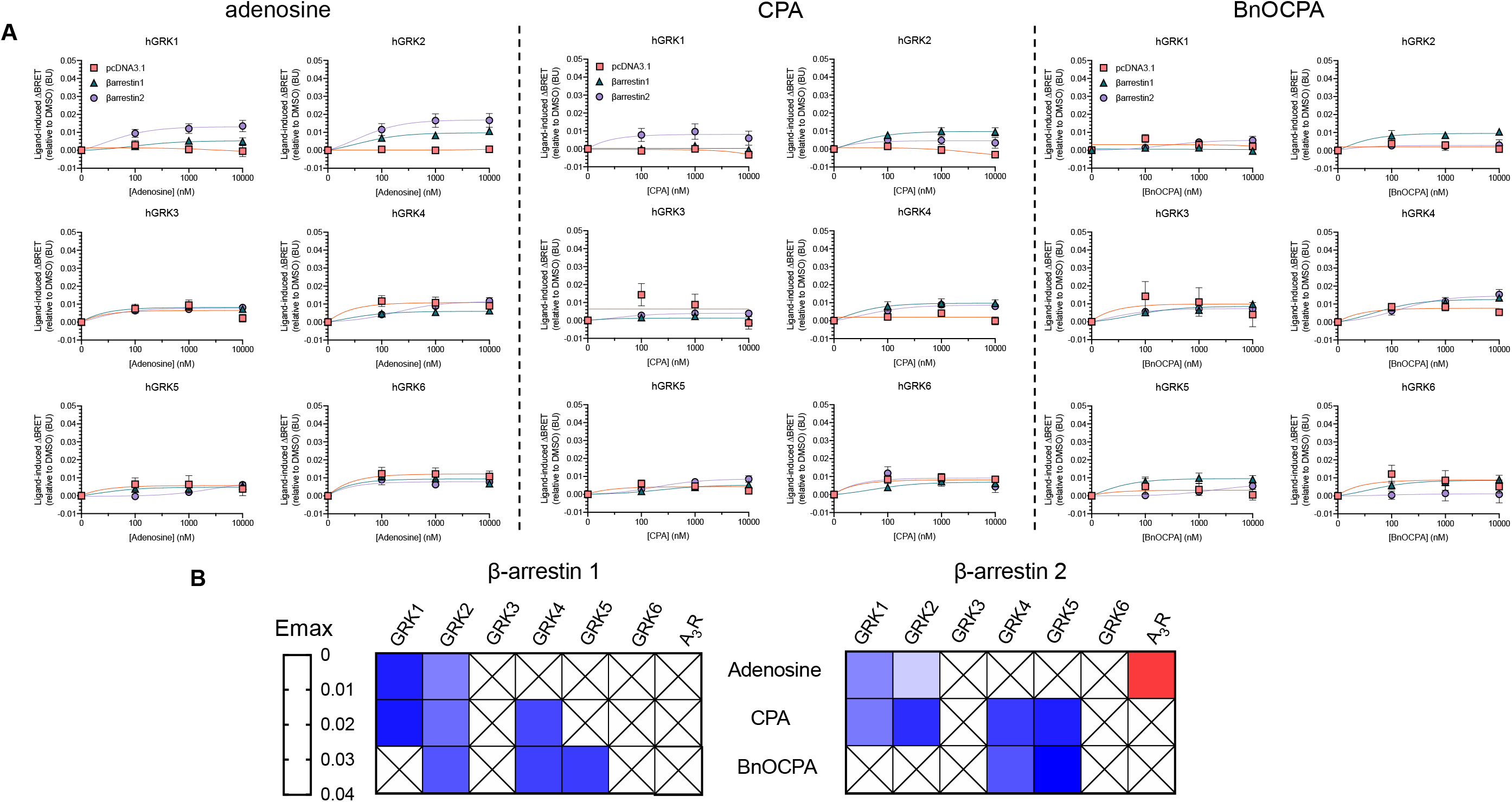
GRK dependence of β-arrestin1 or β-arrestin2 recruitment to the hA_1_R. **A**, Human G protein receptor kinase (hGRK) isoforms 1 – 6 were over expressed (5-fold relative to A_1_R-Nluc), in the presence of control vector (pcDNA3.1) or β-arrestin1-YFP or β-arrestin2-YFP. BRET coupling was examined for each of these combinations for adenosine (left panels); BnOCPA (middle panels) or CPA (right panels). **B**, Heat map describing the peak maximum β-arrestin1 (left panel) and β-arrestin2 (right panel) recruitment for hA_1_R in the presence of the 6 GRK isoforms. A_3_R β-arrestin recruitment is included as a control. In all cases minimal β-arrestin recruitment was observed for the three agonists at the A_1_R.

**Supplementary Fig. 5.**
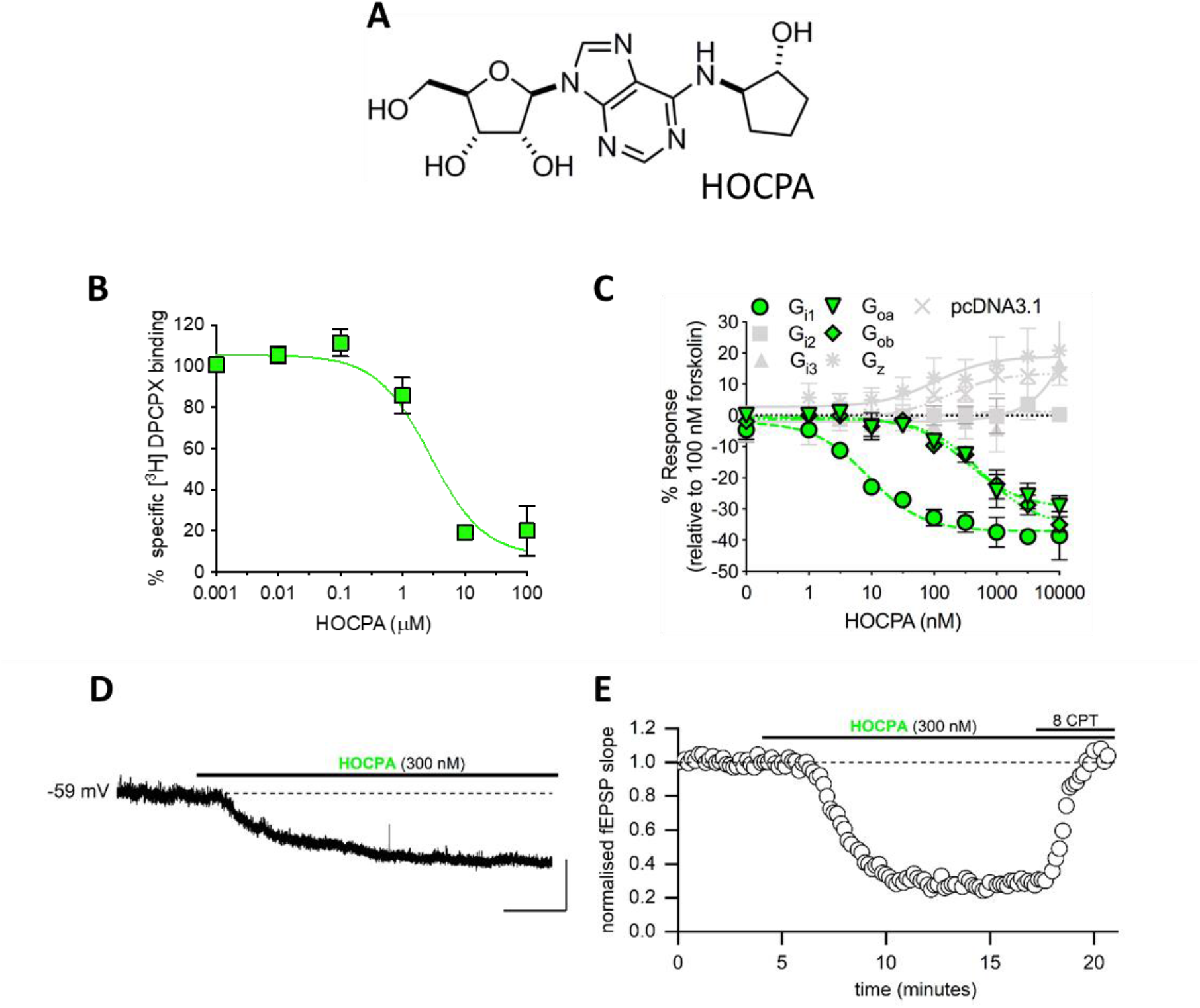
HOCPA does not show Gα selectivity and does not discriminate between pre- and postsynaptic A_1_Rs. **A**, Chemical structure of HOCPA. **B**, Binding of HOCPA was measured via its ability to displace [^3^H]DPCPX from CHO-K1-hA_1_R cells membranes. **C**, The ability of HOCPA to inhibit forskolin-stimulated (100 nM) cAMP production in PTX pre-treated (200 ng/ml) CHO-K1-hA_1_R cells, transfected with PTX-insensitive Gi1, Gi2, Gi3, Goa, Gob, Gz or control (pcDNA3.1). In contrast to BnOCPA, HOCPA shows no selectivity between Goa and Gob. All data are presented as mean ± SEM, of *n* = 4 – 5 individual replicates. **D**, Example membrane potential trace. HOCPA (300 nM) induced hyperpolarisation (mean hyperpolarisation 5.3 ± 0.5 mV, n = 6 cells). Scale bars measure 5 mV and 50 s. **E**, Graph plotting normalised fEPSP slope against time for a single experiment. HOCPA caused a ∼80 % reduction in fEPSP slope, which was reversed by the A_1_R antagonist 8CPT (4 µM). Similar results were observed in 4 slices.

**Supplementary Fig. 6.**
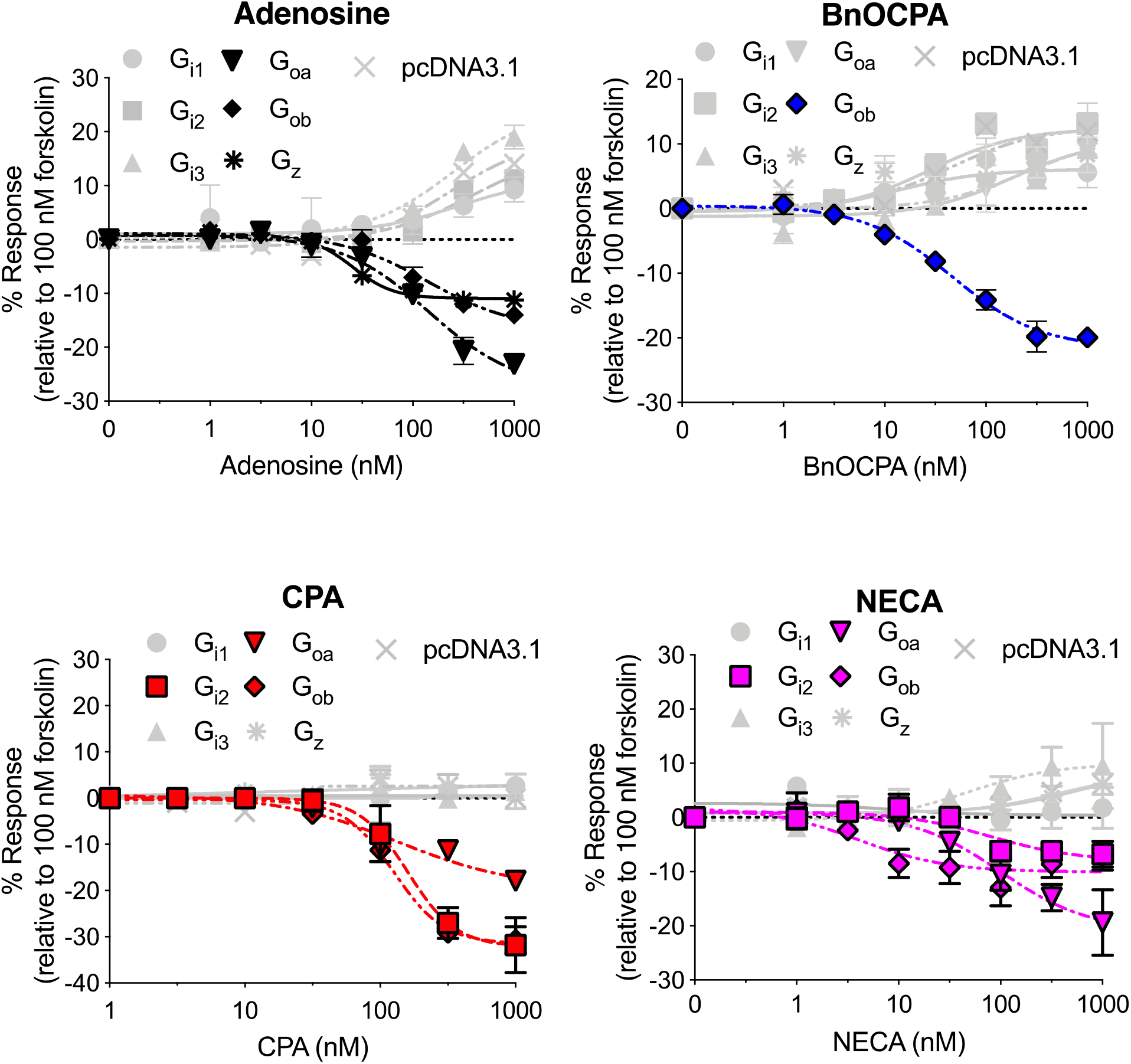
Prototypical and atypical A_1_R agonists display differing Gαi/o activation profiles. The ability of adenosine, BnOCPA, CPA and NECA to activate each individual Gi/o/z subtype was determined in CHO-K1-hA_1_R cells, transfected with PTX-insensitive G proteins or control (pcDNA3.1). cAMP levels were measured following 30 minute co-stimulation with 100 nM forskolin and each agonist. Adenosine displayed an ability to inhibit cAMP production via activation of Gi2, Goa, Gob, and Gz; CPA and NECA via Gi2, Goa and Gob, and BnOCPA exclusively via Gob. Data represented as the average level of cAMP production relative to that observed upon stimulation with 100 nM forskolin, ± SEM, of *n* = 4 – 6 individual replicates. Stimulation of cAMP production reflects activation of endogenous Gs by the A_1_R and is in agreement with previous observations^1-3^.

**Supplementary Fig. 7.**
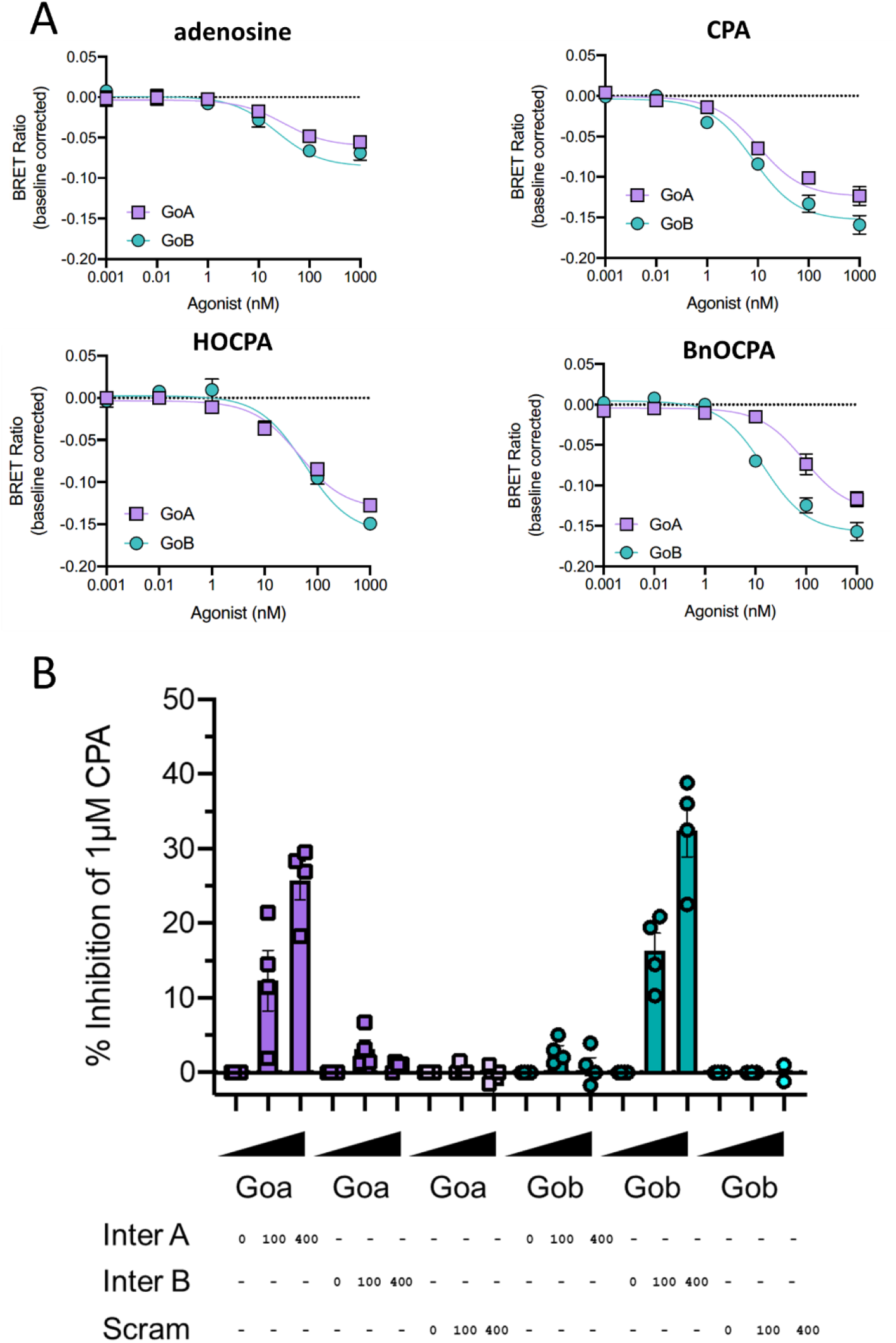
TruPath assays of Goa and Gob activation and the influence of interfering peptides against Goa and Gob. **A** Concentration-response curves (from 6 – 8 biological replicates performed in duplicate) for the agonist-induced dissociation of Gα and Gβγ subunits in the TruPath BRET assay for Goa and Gob activation. Ratios have been baseline corrected with respect to a blank sample. **B** Effects of increasing doses (in ng of plasmid) of interfering and scrambled peptides on the BRET ratio obtained from Goa and Gob in response to 1 μM CPA (4 biological replicates performed in duplicate). Inhibition of the CPA-induced BRET signal is only seen when the interfering peptide is used against its cognate Go isoform. The scrambled Goa peptide has no effect on the CPA-induced BRET signal induced by either Goa or Gob.

**Supplementary Fig. 8.**
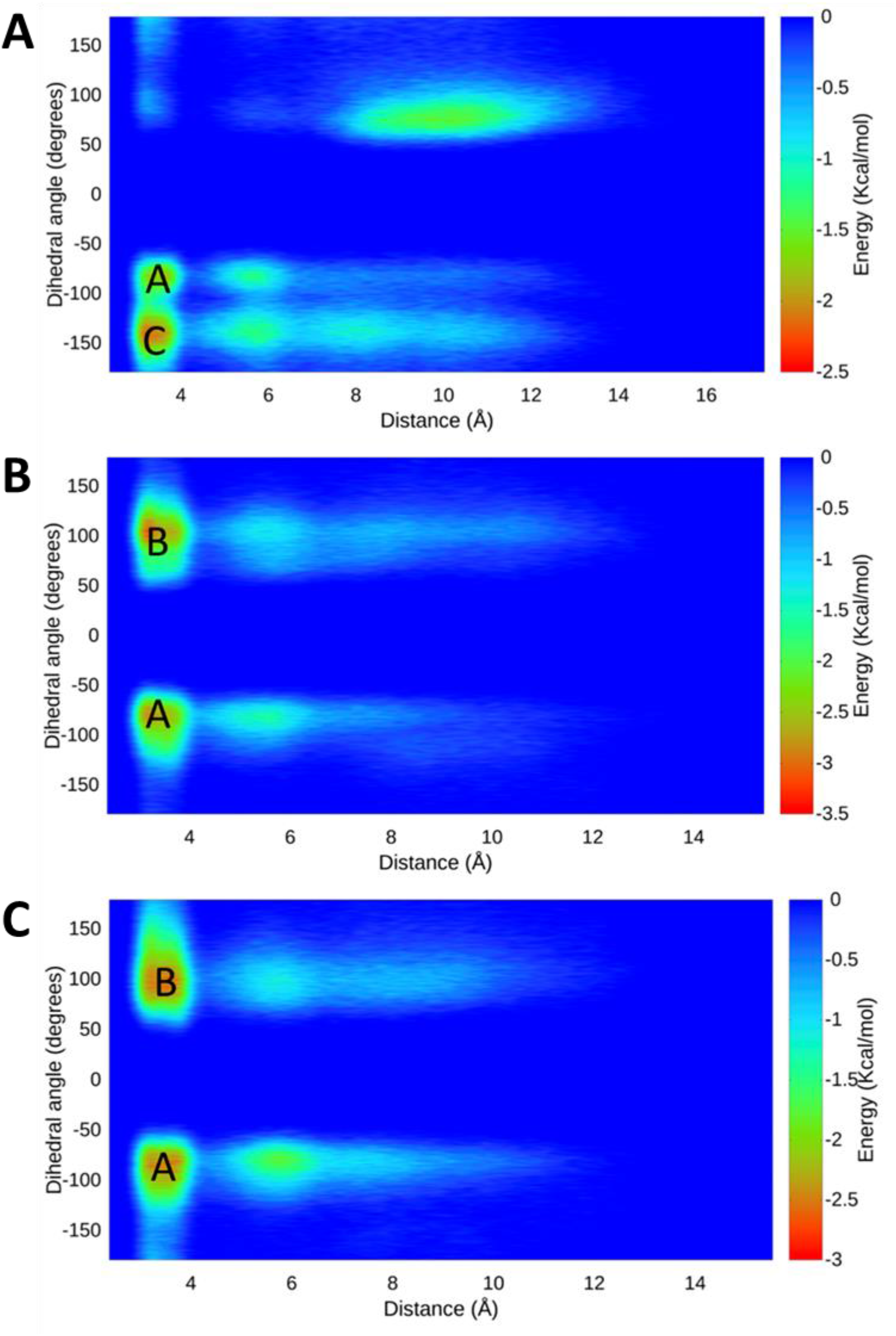
Energy surfaces obtained from metadynamics simulations of BnOCPA. Energy surface obtained by integrating the Gaussian terms deposited during three well-tempered metadynamics replicas (panels **A**, **B** and **C**). X axes report the distance between the E172^ECL2^ carboxyl carbon and the positively charged K265^ECL3^ nitrogen atom; Y axes indicate the dihedral angle formed by the 4 atoms linking BnOCPA cyclopentyl ring to the phenyl moiety. The three energy minima (A, B and C) correspond to the three binding modes proposed for BnOCPA (Modes A, B, C in Fig. 3d to f, respectively).

**Supplementary Fig. 9.**
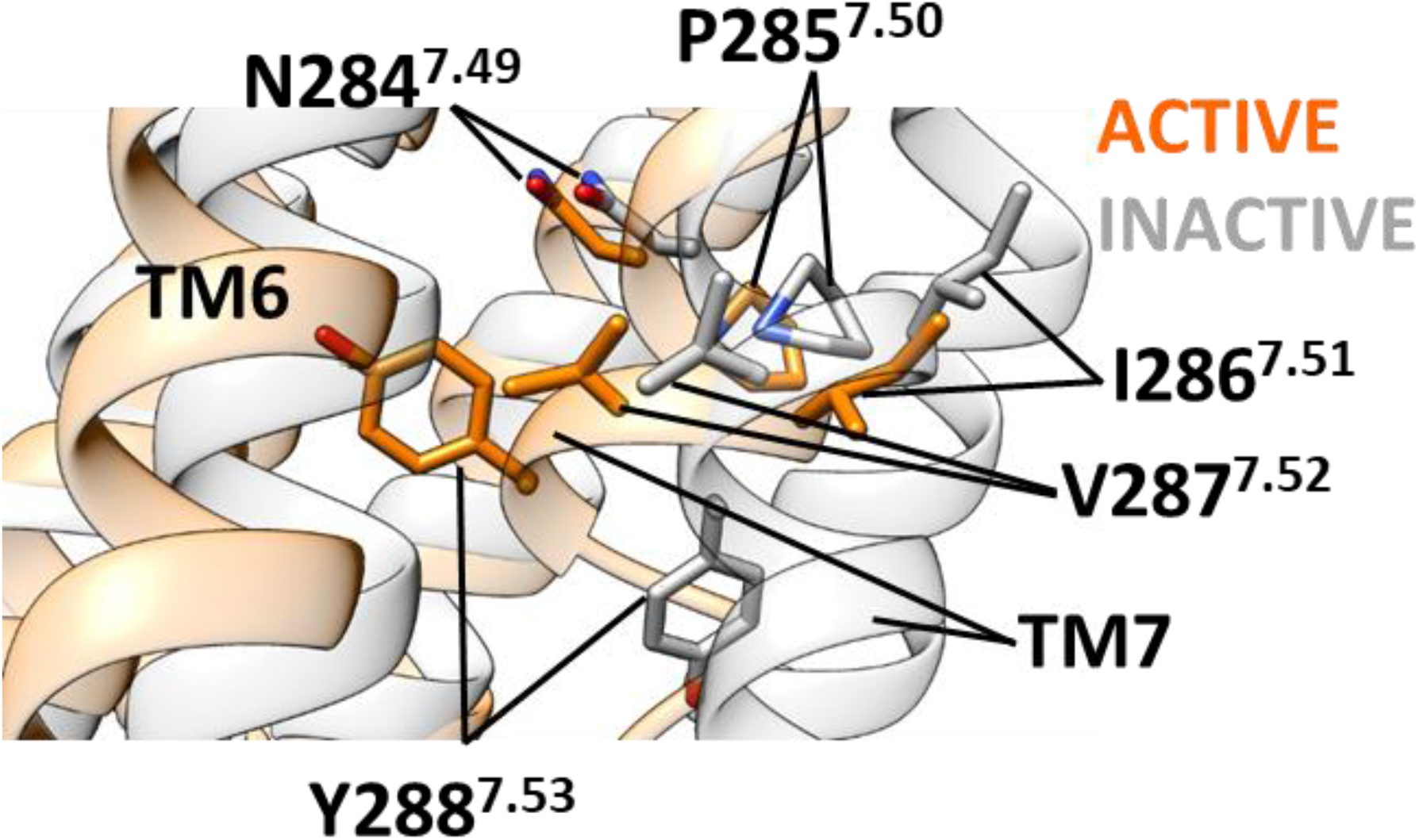
The conserved NPXXY motif (N^7.49^ PIV Y^7.53^) in the A_1_R. The root mean square deviation (RMSD) was computed with respect to the A_1_R inactive conformation. Compared to the inactive conformation (grey), in the active state (orange) the distal portion of TM7 is moved towards the TM bundle core (which is responsible for G protein binding). Starting from the active conformation (orange) and in absence of bound G protein, simulations should allow the structure to partially relax towards the inactive state (grey) with a dynamic influenced by the orthosteric ligand.

**Supplementary Fig. 10.**
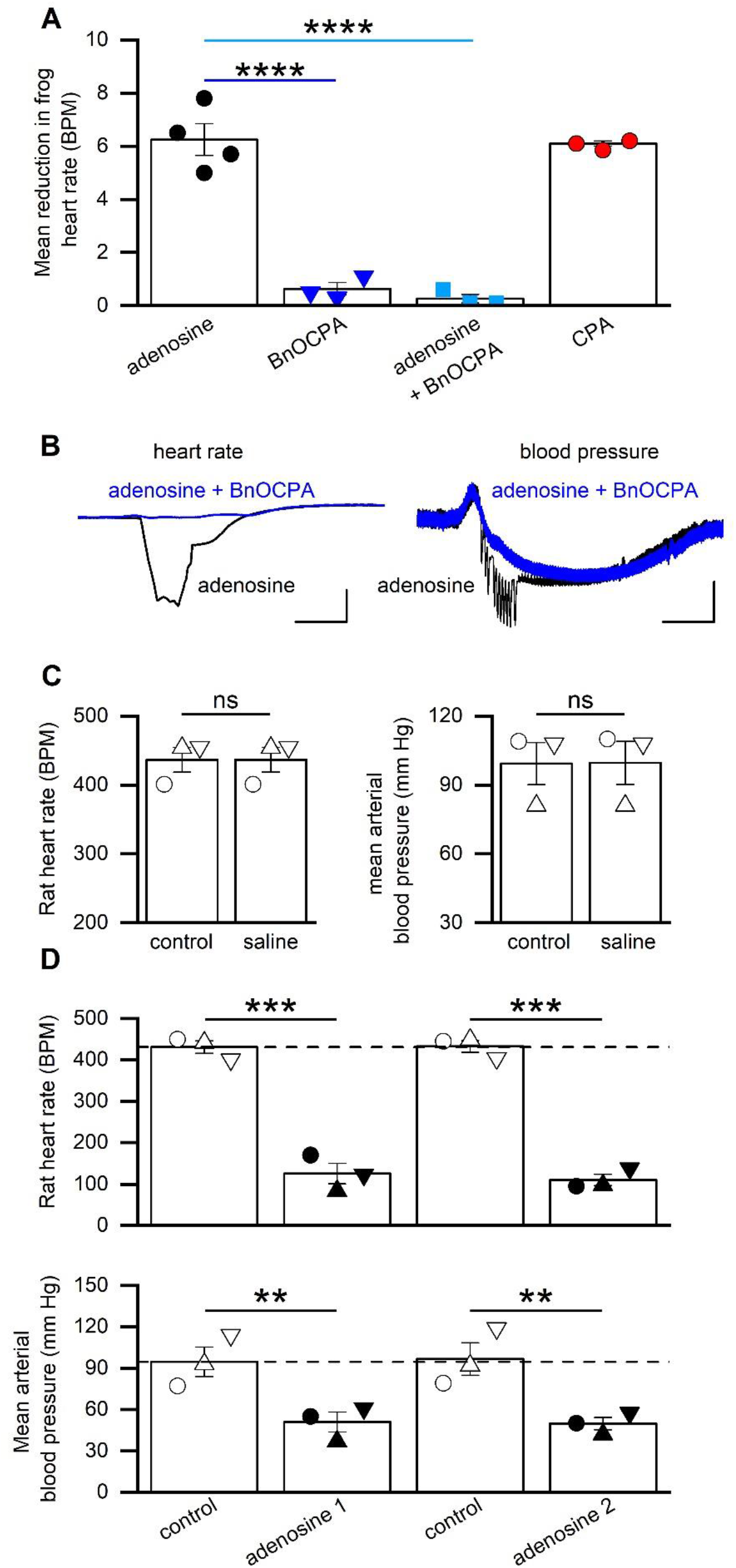
Actions of BnOCPA on frog heart rate and controls for anaesthetised rat experiments. **A**, Data summary for 3 – 4 isolated frog heart preparations. Application of adenosine (30 µM) reduced heart rate (HR) from 41.8 ± 1.3 BPM to 35.5 ± 1.3 BPM. BnOCPA (300 nM) had no effect on HR (42.8 ± 1.2 BPM vs 42.1 ± 1.2 BPM; change 0.6 ± 0.2 BPM), an effect that was significantly different from that of adenosine (blue line; P = 2.22 x 10^-5^). BnOCPA significantly (cyan line; P = 1.31 x 10^-5^) reduced the effects of subsequent adenosine applications (from a reduction of 6.3 ± 0.6 BPM to 0.3 ± 0.2 BPM). CPA (300 nM) reduced HR by 6.1 ± 0.1 BPM, a value similar to that of adenosine. One way ANOVA on the difference in HR across the 4 conditions (F(3,9) = 64.64; P = 2.070 x 10^-6^), with the reported Bonferroni-corrected P values. **B**, Representative traces from a urethane-anaesthetised, spontaneously breathing rat. BnOCPA blocks the effect of adenosine on heart rate (left traces), but only prevents the early phase of adenosine-induced hypotension (right trace). Data taken from the trace in Fig. 5. Scale bars measure 100 BPM or 20 mm Hg and 6 s. **C**, Data summary for 3 urethane-anaesthetised, spontaneously breathing rats. Bar charts showing that injection of 0.9 % saline (equivalent volume to drug experiments) had no effect (paired t-test) on either HR (P = 1) or mean arterial blood pressure (MAP; P = 0.422). **D**, Data summary for 3 urethane-anaesthetised, spontaneously breathing rats. Repeated adenosine injections have the same significant effect on HR (P = 1.40 x 10^-4^ and 1.02 x 10^-4^, respectively) and MAP (P = 0.012 and 0.008, respectively) and thus show no run down. One-way RM ANOVA for both HR (Greenhouse-Geisser corrected F(1.97,3.94) = 96.79, P = 4.48 x 10^-4^, and MAP (F(1.10,2.20) = 19.46, P = 0.040) from 3 animals. In **C** and **D**, each symbol represents data from a single rat. ns, not significant; **, P < 0.02; ***, P < 0.001; ****, P < 0.0001.

**Supplementary Fig. 11.**
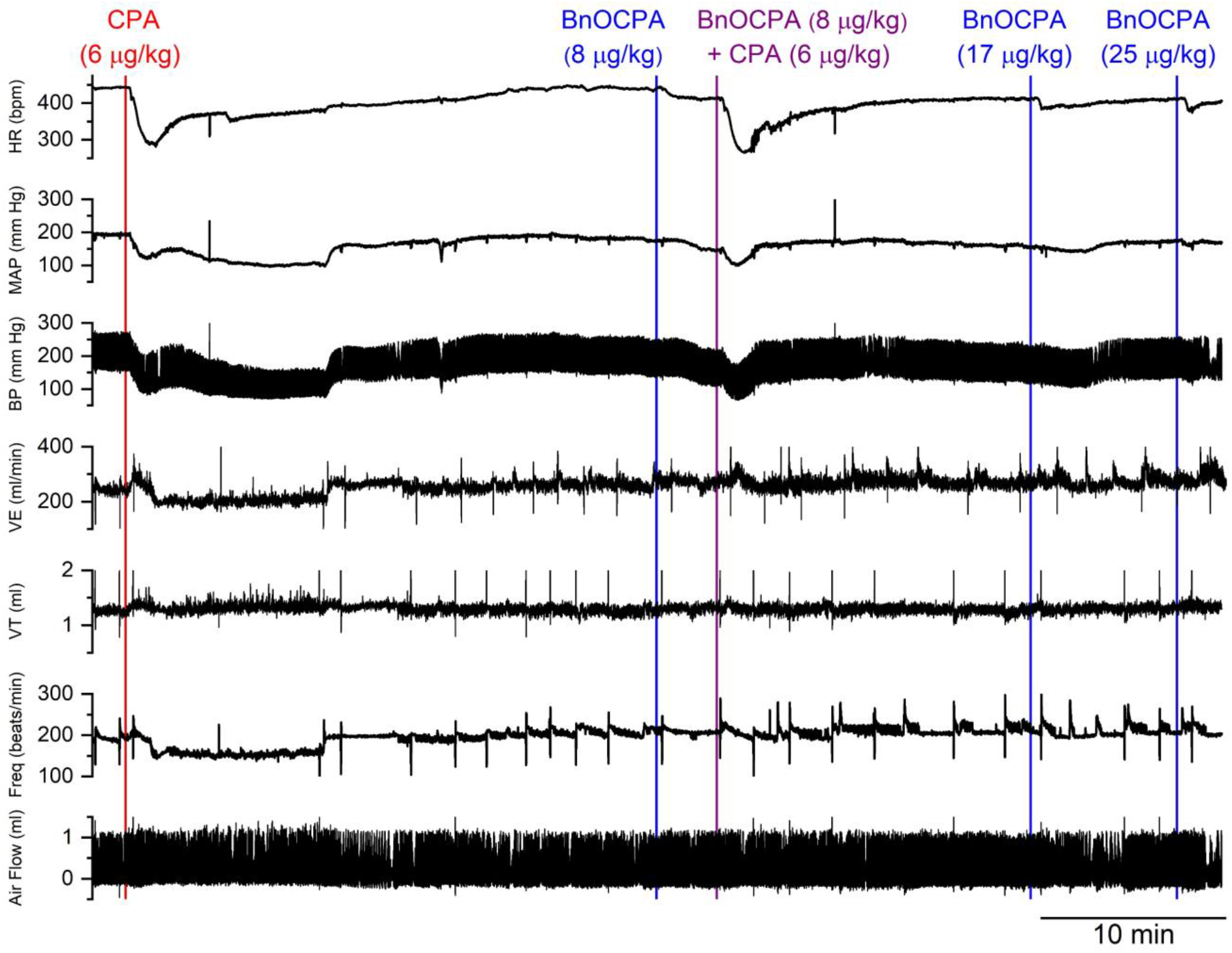
BnOCPA has no actions on cardiorespiratory parameters, but antagonizes the effects of CPA. Examples of traces from a single spontaneously breathing urethane-anaesthetised rat showing: blood pressure (BP), from which heart rate (HR), and mean arterial pressure (MAP) are calculated, and tracheal tube airflow, from which respiratory frequency (Freq), tidal volume (V_T_) and minute ventilation (VE) are calculated. Applications of CPA (6 µg/kg; red vertical line), BnOCPA (8 µg/kg, 17 µg/kg, and 25 µg/kg; blue vertical lines), and the co-application (purple vertical line) of BnOCPA (8 µg/kg) and CPA (6 µg/kg) are shown by the vertical lines. BnOCPA and CPA were given as a 350 µL/kg IV bolus. The intravenous cannula was flushed with 0.9% saline to remove compounds in the tubing between drug applications. The second phase of the blood pressure response following the first dose of CPA is likely the result of the hyponea.

### Supplementary Tables

**Supplementary Table 1.** Binding affinities and efficacies at human A_1_R, A_2_AR and A_3_R expressed in CHO-KI cells.

|  | pK<sub>i</sub><sup>a</sup> | hA<sub>1</sub>R<br>pIC<sub>50</sub><sup>b</sup> | Range<sup>c</sup> | pK<sub>i</sub><sup>d</sup> | hA<sub>2A</sub>R<br>pEC<sub>50</sub><sup>b</sup> | Range<sup>c</sup> | pK<sub>i</sub><sup>d</sup> | hA<sub>3</sub>R<br>pIC<sub>50</sub><sup>b</sup> | Range<sup>c</sup> |
| --- | --- | --- | --- | --- | --- | --- | --- | --- | --- |
| Adenosine | 5.02 ± 0.10 | 8.45 ± 0.2 | 55.7 ± 4.0 | 5.74 ± 0.11 | 6.09 ± 1.3 | 38.1 ± 3.16 | 6.04 ± 0.1 | 8.34 ± 0.2 | 39.8 ± 2.3 |
| CPA | 6.65 ± 0.14*** | 9.26 ± 0.3*** | 48.97 ± 0.7 | N.D | N.D | N.D | <4.0<sup>#</sup> | N.R. | N.R. |
| NECA | 6.45 ± 0.06*** | 9.05 ± 0.2*** | 34.33 ± 4.0 | 6.36 ± 0.09 | 7.94 ± 0.1** | 42.99 ± 4.32 | 6.82 ± 0.12 | 9.2 ± 0.31** | 33.43 ± 3.1 |
| HOCPA | 5.81 ± 0.16*** | 9.08 ± 0.1** | 60.52 ± 1.5 | <4.0<sup>#</sup> | 5.49 ± 0.1* | 41.66 ± 6.1 | 6.23 ± 0.2 | 7.61 ± 0.22*** | 45.0 ± 2.1 |
| BnOCPA | 6.47 ± 0.11*** | 9.17 ± 0.3*** | 49.0 ± 0.66 | <4.0<sup>#</sup> | 5.60 ± 0.01 | 49.0 ± 4.6 | <4.0<sup>#</sup> | N.R. | N.R. |

Average data ± SEM of 3 - 19 individual replicates
<sup>a</sup> Negative logarithm of agonist concentration displacing 50% bound [<sup>3</sup>H]-DPCPX
<sup>b</sup> Negative logarithm of agonist concentration producing half-maximal response
<sup>c</sup> Range of response observed upon agonist stimulation, as a percentage of response obtained upon stimulation with 10 μM forskolin
<sup>d</sup> Negative logarithm of agonist concentration displacing 50% bound
<sup>#</sup> Full estimates of the binding constant were not possible due to failure to generate a full inhibition curve.
N.D. – Not determined, N.R. – No response.
Statistical difference between each agonist and adenosine was calculated using a one-way ANOVA with Dunnett's post-test (\*\* P < 0.01; \*\*\* P < 0.001).

**Supplementary Table 2.** Binding affinities and efficacies at rat A_1_R, A_2_AR and A_3_R expressed in CHO-KI cells.

|  | pK<sub>i</sub><sup>a</sup> | rA<sub>1</sub>R<br>pIC<sub>50</sub><sup>b</sup> | Range<sup>c</sup> | pK<sub>i</sub><sup>a</sup> | rA<sub>2A</sub>R<br>pEC<sub>50</sub><sup>b</sup> | Range<sup>c</sup> | pK<sub>i</sub><sup>d</sup> | rA<sub>3</sub>R<br>pIC<sub>50</sub><sup>b</sup> | Range<sup>c</sup> |
| --- | --- | --- | --- | --- | --- | --- | --- | --- | --- |
| **Adenosine** | 5.41 ± 0.18 | 7.63 ± 0.1 | 34.40 ± 1.04 | 5.74 ± 0.11 | 7.58 ± 0.18 | 37.78 ± 2.76 | 5.89 ± 0.09 | 7.17 ± 0.18 | 66.05 ± 5.4 |
| **CPA** | 6.80 ± 0.14*** | 9.47 ± 0.16*** | 36.31 ± 2.57 | <4.0<sup>#</sup> | 5.55 ± 0.19*** | 39.45 ± 3.47 | N.D | 7.41 ± 0.13 | 68.79 ± 3.67 |
| **NECA** | 6.32 ± 0.13*** | 8.65 ± 0.77** | 37.17 ± 5.49 | 6.36 ± 0.09 | 8.37 ± 0.18* | 37.78 ± 2.65 | 6.43 ± 0.11** | 8.81 ± 0.18** | 63.02 ± 5.14 |
| **HOCPA** | 6.27 ± 0.14*** | 9.01 ± 0.01*** | 34.40 ± 1.59 | 4.86 ± 0.12** | 5.69 ± 0.20*** | 48.61 ± 6.42* | 6.17 ± 0.02* | 7.41 ± 0.12 | 68.79 ± 3.70 |
| **BnOCPA** | 6.25 ± 0.16** | 8.92 ± 0.14*** | 39.45 ± 1.11 | 5.03 ± 0.11** | 5.00 ± 0.13*** | 37.15 ± 2.76 | 5.27 ± 0.12 | 6.73 ± 0.13* | 65.16 ± 3.80 |

Average data ± SEM of 3 - 19 individual replicates
<sup>a</sup> Negative logarithm of agonist concentration displacing 50% bound CA200645
<sup>b</sup> Negative logarithm of agonist concentration producing half-maximal response
<sup>c</sup> Range of response observed upon agonist stimulation, as a percentage of response obtained upon stimulation with 10 μM forskolin
<sup>d</sup> Negative logarithm of agonist concentration displacing 50% bound AV039
<sup>#</sup> Full estimates of the binding constant were not possible due to failure to generate a full inhibition curve.
N.D. – Not determined.
Statistical difference between each agonist and adenosine was calculated using a one-way ANOVA with Dunnett's post-test (\* P < 0.05; \*\* P < 0.01; \*\*\* P < 0.001).

**Supplementary Table 3.** Transient hydrogen bonds between α4-β6 loop residue 317 (N317 in Goa, H317 in Gob), the α3-β5 loop residue D263, and the residue on H8 of the A_1_R (Ballesteros Weinstein enumeration in superscript).

| **A<sub>1</sub>R - Gα Interactions** |  |  |  |  |  |
| --- | --- | --- | --- | --- | --- |
|  | **Coupling Systems** |  |  | **Non-coupling Systems** |  |
|  | **Occupancy (%frames)** |  |  | **Occupancy (%frames)** |  |
| **A<sub>1</sub>R - Gα hydrogen bond** | **BnOCPA mode D:Gob** | **BnOCPA mode B:Gob** | **HOCPA:Gob** | **BnOCPA mode D:Goa** | **BnOCPA mode B:Goa** |
| **H317-Q293<sup>8,48</sup>** | **1.7** | **0.5** | **2.8** | **6.9** | **10.3** |
| **D263-Q293<sup>8,48</sup>** | **0.4** | **0.4** | **1.5** | **9.2** | **0.6** |
| **K294<sup>8,49</sup>-D263** | **0.0** | **0.0** | **0.1** | **4.0** | **2.9** |
| **R296<sup>8,51</sup>-D263** | **0.1** | **0.5** | **0.0** | **10.7** | **0.0** |

### Supplementary Movies

**Supplementary Movie 1**

**Molecular dynamics dynamic docking simulation of BnOCPA binding to the apo A_1_R** Extracellular (left) and orthogonal (right) views of BnOCPA (stick and transparent sphere representation) simulation of binding to the apo A_1_R (white ribbon). Protein residues within 4 Å from the ligand atoms are shown (stick representation). Hydrogen bonds are highlighted as red dotted lines. Soon after it reached the orthosteric site, BnOCPA engaged N254^6.55^ in a bi-dentate hydrogen bond. The ribose moiety, initially involved in an intramolecular hydrogen bond with the purine ring, interacts with side chains of internal residues, such as the key residue for receptor activation, T277^7.42^. The benzyl moiety initially explores binding mode A, then moves to mode B (after about 720 ns).

**Supplementary Movie 2**

**Molecular dynamics dynamic docking simulation of HOCPA binding to the apo A_1_R** Extracellular (left) and orthogonal (right) views of HOCPA (stick and transparent sphere representation) simulated binding to the apo A_1_R (white ribbon). Protein residues within 4 Å from the ligand atoms are shown (stick representation). Hydrogen bonds are highlighted as red dotted lines. Soon after it entered into the orthosteric site, HOCPA engaged N254^6.55^ in a bi-dentate hydrogen bond. In analogy to BnOCPA (Extended Data Movie 2) the ribose moiety, initially involved in an intramolecular hydrogen bond with the purine ring, interacts with side chains of inner located residues, such as the key residue for receptor activation T277^7.42^. Further hydrogen bonds are formed between the cyclopentyl hydroxyl group and the ionic bridge between E172^ECL2^ and K265^ECL3^.

**Supplementary Movie 3**

**Molecular dynamics dynamic docking simulation of the Gob GαCT to the BnOCPA:A_1_R complex.** Intracellular view of the A_1_R (white ribbon and transparent surface) during the binding simulations of Gob GαCT (also denoted as H5 – black ribbon). The transparent ribbon shows the position of the Gi2 GαCT as reported in the cryo-EM A_1_R structure 6D9H. The supervision algorithm is switched off after about 43 ns of productive simulation.

Supplementary Movie 4

**Molecular dynamics simulation of the BnOCPA:A_1_R:Goa(α subunit) complex.** Intracellular view of the A_1_R (white ribbon and cyan stick representation) bound to the Goa α subunit (orange ribbon and green stick representation) during one MD replica. After about 300 ns of simulation the system undergo a conformational transition characterized by transient hydrogen bonds between the receptor H8 (Q293^8.48^ and R296^8.51^) and the Goa residues located on the α3-β5 (D263) and α4-β6 (N317) loops.

**Supplementary Movie 5**

**Molecular dynamics simulation of the BnOCPA:A_1_R:Gob(α subunit) complex.** Intracellular view of the A_1_R (white ribbon and cyan stick representation) bound to the Gob α subunit (orange ribbon and green stick representation) during one MD replica. The system shows lower flexibility than BnOCPA:A_1_R:Goa. Stable interactions between the Gob α3-β5 loop and the α5 (GαCT) positively charged K347 and R350 occurs.

### Supplemental Data Files 1

**BnOCPA pharmacokinetics Excel spreadsheet**

## Methods

### Materials and Methods

#### Approvals

All experiments involving animals were conducted with the knowledge and approval of the University of Warwick Animal Welfare and Ethical Review Board, and in accordance with the U.K. Animals (Scientific Procedures) Act (1986) and the EU Directive 2010/63/EU. *In vivo* cardiorespiratory studies were conducted under the auspices of UK PPL 70/8936 and the chronic neuropathic pain studies under the auspices of P9D9428A9.

#### Preparation of hippocampal slices

Sagittal slices of hippocampus (300-400 µm) were prepared from male Sprague Dawley rats, at postnatal days 12-20^1^. Rats were kept on a 12-hour light-dark cycle with slices made 90 minutes after entering the light cycle. In accordance with the U.K. Animals (Scientific Procedures) Act (1986), rats were killed by cervical dislocation and then decapitated. The brain was removed, cut down the midline and the two sides of the brain stuck down to a metal base plate using cyanoacrylate glue. Slices were cut along the midline with a Microm HM 650V microslicer in cold (2-4°C), high Mg^2+^, low Ca^2+^ artificial cerebrospinal fluid (aCSF), composed of (mM): 127 NaCl, 1.9 KCl, 8 MgCl_2_, 0.5 CaCl_2_, 1.2 KH_2_PO_4_, 26 NaHCO_3_, 10 D-glucose (pH 7.4 when bubbled with 95% O_2_ and 5% CO_2_, 300 mOSM). Slices were stored at 34°C for 1-6 hours in aCSF (1 mM MgCl_2_, 2 mM CaCl_2_) before use.

#### Extracellular recording

A slice was transferred to the recording chamber, submerged in aCSF and perfused at 4-6 ml·min^-1^ (32 ± 0.5°C). The slice was placed on a grid allowing perfusion above and below the tissue and all tubing was gastight (to prevent loss of oxygen). An aCSF-filled glass microelectrode was placed within stratum radiatum in area CA1 and recordings were made using either a differential model 3000 amplifier (AM systems, WA USA) or a DP-301 differential amplifier (Warner Instruments, Hampden, CT USA). Field excitatory postsynaptic potentials (fEPSPs) were evoked with either an isolated pulse stimulator model 2100 (AM Systems, WA) or ISO-Flex (AMPI, Jerusalem, Israel). For fEPSPs a 10-20 minute baseline was recorded at a stimulus intensity that gave 40-50% of the maximal response. Signals were acquired at 10 kHz, filtered at 3 kHz and digitised on line (10 kHz) with a Micro CED (Mark 2) interface controlled by Spike software (Vs 6.1, Cambridge Electronic Design, Cambridge UK) or with WinLTP^2^. For fEPSP slope, a 1 ms linear region after the fibre volley was measured. Extracellular recordings were made independently on two electrophysiology rigs. As the data obtained from each rig was comparable, both sets of data have been pooled.

#### Seizure model

Seizure activity was induced in hippocampal slices using nominally Mg^2+^-free aCSF that contained no added Mg^2+^ and with the total K^+^ concentration increased to 6 mM with KCl. Removal of extracellular Mg^2+^ facilitates depolarisation via glutamate N-methyl-D-aspartate (NMDA) receptor activation. Increasing the extracellular concentration of K^+^ depolarises neurons leading to firing and release of glutamate to sustain activity. Both the increase in K^+^ concentration and removal of Mg^2+^ are required to produce spontaneous activity in hippocampal slices^3^. Spontaneous activity was measured with an aCSF-filled microelectrode placed within stratum radiatum in area CA1.

#### Whole-cell patch clamp recording from hippocampal pyramidal cells

A slice was transferred to the recording chamber and perfused at 3 ml·min^-1^ with aCSF at 32 ± 0.5°C. Slices were visualized using IR-DIC optics with an Olympus BX151W microscope (Scientifica) and a CCD camera (Hitachi). Whole-cell current- and voltage-clamp recordings were made from pyramidal cells in area CA1 of the hippocampus using patch pipettes (5–10 MΩ) manufactured from thick walled glass (Harvard Apparatus, Edenbridge UK) and containing (mM): potassium gluconate 135, NaCl 7, HEPES 10, EGTA 0.5, phosphocreatine 10, MgATP 2, NaGTP 0.3 and biocytin 1 mg ml^−1^ (290 mOSM, pH 7.2). Voltage and current recordings were obtained using an Axon Multiclamp 700B amplifier (Molecular Devices, USA) and digitised at 20 KHz. Data acquisition and analysis was performed using Pclamp 10 (Molecular Devices, USA). For voltage clamp experiments, CA1 pyramidal cells were held at -60 mV. Peptides to interfere with G protein signalling were introduced via the patch pipette into the recorded cell. The cell was held for at least 10 minutes before adenosine (10 µM) was added to induce an outward current.

#### Frog heart preparation

Young adult male *Xenopus leavis* frogs were obtained from Portsmouth Xenopus Resource Centre. Frogs were euthanized with MS222 (0.2 % at a pH of 7), decapitated and pithed. The animals were dissected to reveal the heart and the pericardium was carefully removed. Heart contractions were measured with a force transducer (AD instruments). Heart rate was acquired via a PowerLab 26T (AD instruments) controlled by LabChart 7 (AD instruments). The heart was regularly washed with Ringer solution and drugs were applied directly to the heart.

#### *In vivo* anaesthetised rat preparation for cardiorespiratory recordings

Anaesthesia was induced in adult male Sprague Dawley rats (230-330 g) with isofluorane (2-4%; Piramal Healthcare). The femoral vein was catheterised for drug delivery. Anaesthesia was maintained with urethane (1.2-1.7 g·kg^-1^; Sigma) in sterile saline delivered via the femoral vein catheter. Body temperature was maintained at 36.7°C via a thermocoupled heating pad (TCAT 2-LV; Physitemp). The trachea was cannulated and the femoral artery catheterised, and both were connected to pressure transducers (Digitimer) to record respiratory airflow and arterial blood pressure, respectively. Blood pressure and airflow signals were amplified using the NeuroLog system (Digitimer) connected to a micro1401 interface and acquired on a computer using Spike2 software (Cambridge Electronic Design). Arterial blood pressure recordings were used to derive heart rate (HR: beats.minute^-1^; BPM), and to calculate mean arterial blood pressure (MAP: Diastolic pressure + ⅓*[Systolic Pressure - Diastolic pressure]). Airflow measurements were used to calculate: tidal volume (V_T_; mL; pressure sensors were calibrated with a 3 mL syringe), and respiratory frequency (*f*; breaths·min^-1^; BrPM). Minute ventilation (VE; mL·min^-1^) was calculated as *f* x V_T_.

Cardiovascular and respiratory parameters were allowed to stabilise before experiments began. A_1_R agonists were administered by intravenous (IV) injection and the changes in HR, MAP, *f*, V_T_, and V_E_ were measured. In pilot studies, the optimal dose of adenosine was determined by increasing the dose until robust and reliable changes in HR and MAP were produced (1 mg·kg^-1^). The dose of CPA was adjusted until equivalent effects to adenosine were produced on HR and MAP (6 µg·kg^-1^). For BnOCPA we initially used 1 µg·kg^-1^, but saw no agonist effect on HR and MAP. To ensure this was not a false negative we increased the dose of BnOCPA (8 µg·kg^-1^), which still gave no agonist effect on HR and MAP. However, as BnOCPA produced an antagonistic effect when co-administered with adenosine (Fig. 5, Supplementary Fig. 10b) and CPA (Fig. 6, Supplementary Fig. 11), it must have reached A_1_Rs at a high enough concentration to be physiologically active. These observations confirmed that the lack of agonistic effects on HR and MAP were not due to a type II error. 8 µg·kg^-1^ BnOCPA was used for all further experiments. All injections were administered IV as a 350 µl·kg^-1^ bolus.

In the experimental studies, rats either:

(1) received an injection of adenosine. After cardiorespiratory parameters returned to baseline (5-10 minutes), rats were given BnOCPA. After allowing sufficient time for any effect of BnOCPA to be observed, rats received adenosine with BnOCPA co-administered in a single injection. After cardiorespiratory parameters returned to baseline, rats were injected with CPA, or
(2) received an injection of CPA. After cardiorespiratory parameters returned to baseline (5-10 minutes) rats were given 8 µg·kg^-1^ BnOCPA. After allowing sufficient time for any effect of BnOCPA to be observed, rats received CPA with 8 µg·kg^-1^ BnOCPA co-administered in a single injection. After cardiorespiratory parameters returned to baseline, rats were injected with successive injections of 17 µg·kg^-1^ and 25 µg·kg^-1^ BnOCPA, with sufficient time given for any effect of BnOCPA to be observed.

To check that the volume of solution injected with each drug did not itself induce a baroreflex response leading to spurious changes in cardiorespiratory responses, equivalent volumes of saline (0.9 %) were injected. These had no effect on either heart rate or MAP (Supplementary Fig. 10c). To confirm that repeated doses of adenosine produced the same response and that the responses did not run-down, rats were given two injections of adenosine (1 mg·kg^-1^). There was no significant difference in the changes in cardiovascular parameters produced by each adenosine injection (Supplementary Fig. 10d).

An additional series of experiments (n = 4) were undertaken to directly compare BnOCPA and CPA on respiration. Adult male Sprague Dawley rats (400-500 g) were anaesthetised with urethane and instrumented as described above, with the exception that the arterial cannulation was not performed.

After allowing the animal to stabilise following surgery, BnOCPA (8 µg·kg^-1^) was administered. After a 20 minutes recovery period CPA (6 µg·kg^-1^) was administered. All injections were administered IV as a 350 µl·kg^-1^ bolus. Changes in *f*, V_T_, and V_E_ were measured. If the dosing occurred close to a respiratory event such as a sigh a second IV dose was administered, with 20 minute recovery periods either side of the injection. Measurements for the effect of BnOCPA were time-matched to when CPA induced a change in respiration in the same preparation. As no difference was observed between the respiratory responses to BnOCPA in these rats (n = 4) and those instrumented for both cardiovascular and respiratory recordings (n = 4), the data were pooled (n = 8; Fig. 6a to d).

#### Spinal nerve ligation (Chung model^4^)

Adult male Sprague-Dawley rats, 7-8 weeks old, weighing around 250 g at the time of Chung model surgery, were purchased from Charles River UK Ltd. The animals were housed in groups of 4 in an air-conditioned room on a 12-hour light/dark cycle. Food and water were available *ad libitum*. They were allowed to acclimatise to the experimental environment for three days by leaving them on a raised metal mesh for at least 40 min. The baseline paw withdrawal threshold (PWT) was examined using a series of graduated von Frey hairs (see below) for 3 consecutive days before surgery and re-assessed on the 6^th^ to 8^th^ day after surgery and on the 13^th^ to 17^th^ day after surgery before drug dosing.

Prior to surgery each rat was anaesthetized with 3% isoflurane mixed with oxygen (2 L·min^-1^) followed by an intramuscular injection of ketamine (60 mg·kg^-1^) plus xylazine (10 mg·kg^-1^). The back was shaved and sterilized with povidone-iodine. The animal was placed in a prone position and a para-medial incision was made on the skin covering the L4-6 level. The L5 spinal nerve was carefully isolated and tightly ligated with 6/0 silk suture. The wound was then closed in layers after a complete hemostasis. A single dose of antibiotics (Amoxipen, 15 mg/rat, intraperitoneally, IP) was routinely given for prevention of infection after surgery. The animals were placed in a temperature-controlled recovery chamber until fully awake before being returned to their home cages. The vehicle (normal saline or DMSO) was administered via the IV route at 1 ml·kg^-1^ and via the intrathecal (IT) route at 10 µl for each injection. The A_1_R-selective antagonist DPCPX (1 mg kg^-1^) and the A_3_R-selective antagonist MRS 1523 (2 mg kg^-1^) were delivered IP 30 mins before vehicle or BnOCPA treatment. The rats with validated neuropathic pain state were randomly divided into 11 groups: vehicle IV, BnOCPA at 1, 3, 6, 10 µg·kg^-1^ IV; vehicle IT, BnOCPA at 0.3, 1, 3 nmol IT; 6 µg·kg^-1^ BnOCPA IV plus 1 mg·kg^-1^ DPCPX IP; 6 µg·kg^-1^ BnOCPA IV plus 2 mg·kg^-1^ MRS1523 IP groups and tested blind to treatment.

To test for mechanical allodynia the animals were placed in individual Perspex boxes on a raised metal mesh for at least 40 minutes before the test. Starting from the filament of lower force, each filament was applied perpendicularly to the centre of the ventral surface of the paw until slightly bent for 6 seconds. If the animal withdrew or lifted the paw upon stimulation, then a hair with force immediately lower than that tested was used. If no response was observed, then a hair with force immediately higher was tested. The highest value was set at 15 g. The lowest amount of force required to induce reliable responses (positive in 3 out of 5 trials) was recorded as the value of PWT. On the testing day, PWT were assessed before and 1, 2 and 4 hours following BnOCPA or vehicle administration. The animals were returned to their home cages to rest (about 30 min) between two neighbouring testing time points. At the end of each experiment, the animals were deeply anaesthetised with isoflurane and killed by decapitation.

#### Rotarod test for motor function

A rotarod test was used to assess motor coordination following intravenous and intraperitoneal administration of BnOCPA. An accelerating rotarod (Ugo Basile) was set so speed increased from 6 to 80 rpm over 170 seconds. Male Sprague Dawley rats (n = 24), 7 weeks of age (212-258g) were trained on the rotarod twice daily for two days (≥2 trials per session) until performance times were stable. On the day of the experiment, three baseline trials were recorded. The compound was administered IP (10 µg/kg, n = 6) or IV via tail vein injection (10 and 25 µg/kg, n = 6 per group). The control group received subcutaneous saline and the positive control group received subcutaneous morphine (15 mg/kg). Latency to fall (seconds) was measured in triplicate at 1, 2, 3 and 5 hours post drug administration. Rotarod studies were approved by the Monash University Animal Ethics Committee in accordance with the Australian Code for the Care and Use of Animals for Scientific Purposes (2013) under Monash AEC protocol number 13333.

#### Constructs, transfections and generation of stable cell lines

To investigate the signalling properties of the rat A_3_R (rA_3_R) and mutants of the human A_1_R (hA_1_R), stable cell lines were generated using Flp-In-CHO cells. Untagged hA_1_R from sigNanoLuciferase (Nluc)-A_1_R in pcDNA3.1+ and untagged rA_3_R from sigNluc-A_3_R in pcDNA3.1+ (both gifted by Dr Steve Briddon (University of Nottingham)) were cloned into the pcDNA5/FRT expression vector (Thermo Fisher Scientific). Mutations within the hA_1_R were made using the QuikChange Lightening Site-Directed Mutagenesis Kit (Agilent Technologies) in accordance with the manufacturer’s instructions. Constructs for generating Goa/b interfering and scrambled peptides were generated by PCR and cloned into the B*amH*I/H*ind*III site of pcDNA3.1- as described in Gilchrist et *al*.,^5^. Prior to the initiator codon a Kozak sequence was included for enhanced translation. The peptide sequences used were: for G_oa_ MGIANNLRGCGLY, for G_ob_ MGIAKNLRGCGLY, and for the scrambled peptide MGLNRGNAYLCIGMG was used. Constructs were sequenced to confirm fidelity. Flp-In-CHO (Thermo Fisher Scientific) cells were generated through co-transfection of the cell line with pcDNA5/FRT expression vector (Thermo Fisher Scientific) containing the WT or mutant hA_1_R, or rA_3_R, and the Flp recombinase expressing plasmid, pOG44 (Thermo Fisher Scientific), in accordance with the manufacturer’s instructions. Co-transfection of cells in a T25 flask, with a total of 5 μg of adenosine receptor (AR)/pcDNA5/FRT and pOG44 (AR:pOG44 ratio of 1:9), was performed using Fugene HD (Promega), at a ratio of 3:1 (v/w) (Fugene:DNA). 24 hours after transfection, cells were harvested and resuspended in growth media containing 600 μg/ml Hygromycin B (Thermo Fisher Scientific), and subsequently seeded into a fresh T25 flask. Media was replaced every 2-3 days and cells stably expressing the receptor of interest were selected using 600 μg/ml Hygromycin B. To generate CHO-K1 cells stably expressing the rat A_2_AR (CHO-K1-rA_2_AR), CHO-K1 cells were seeded onto a 6-well plate and transfected with 1 μg rA_2_AR using Fugene HD (Promega) at a ratio of 3:1 (v/w) (Fugene:DNA). 48 hours after transfection, media was replaced with growth media containing 800 μg/ml G418 (Thermo Fisher Scientific) and changed every 2-3 days until cells were >80% confluent. To investigate rat A_1_R-mediated signalling, CHO-K1 cells seeded onto a 6-well plate were transiently transfected with 1 μg rat A_1_R (rA_1_R) using Fugene HD (Promega) at a ratio of 3:1 (v/w) (Fugene:DNA), for 48 hours. The plasmids encoding the rA_1_R and rA_2_AR (Nluc-A_1_R/pcDNA3.1(+) and Nluc-A_2_AR/pcDNA3.1(+), respectively) were kindly gifted by Stephen Hill and Stephen Briddon (University of Nottingham).

#### Cell signaling assays

CHO cell lines expressing ARs of interest (including mutants of the hA_1_R) were routinely cultured in Ham’s F12 nutrient mix supplemented with 10% foetal bovine serum (FBS), at 37°C with 5% CO_2_, in a humidified atmosphere. For cAMP accumulation experiments, cells were seeded at a density of 2000 cells per well of a white 384-well optiplate and stimulated, for 30 minutes, with a range of agonist concentrations (100 pM – 100 μM) in the presence of 25 μM rolipram (Cayman Chemicals). For cAMP inhibition experiments, cells were co-stimulated with 1 μM forskolin and a range of agonist concentrations (1 pM – 100 μM), in the presence or absence of 1 μM antagonist. cAMP levels were then determined using a LANCE® cAMP kit as described previously^6, 7^.

For determination of individual Gαi/o/z couplings, CHO-K1-hA_1_R cells (made in house) were transfected with pcDNA3.1-GNAZ or, pcDNA3.1 containing pertussis toxin (PTX) insensitive Gαi/o protein mutants (C351I, C352I, C351I, C351I, C351I, for G_i1_, G_i2_, G_i3_, Goa, Gob, respectively, obtained from cDNA Resource Center; www.cdna.org), using 500 ng plasmid and Fugene HD at a 3:1 (v/w) (Fugene:Plasmid) ratio. Cells were then incubated for 24 hours before addition of 100 ng/ml PTX, to inhibit activity of endogenous Gα_i/o_, and then incubated for a further 16-18 hours. Transfected cells were then assayed as per cAMP inhibition experiments, but co-stimulated with agonist and 100 nM forskolin.

#### β-arrestin recruitment assays

HEK 293T cells were routinely grown in DMEM/F-12 GlutaMAX^TM^ (Thermo Fisher Scientific) supplemented with 10% foetal bovine serum (FBS; F9665, Sigma-Aldrich) and 1x antibiotic-antimycotic solution (DMEM complete; Thermo Fisher Scientific). For analysis of β-arrestin recruitment following ligand stimulation at the hA_1_R or hA_3_R, HEK 293T cells in a single well of 6-well plate (confluency ≥80%) were transiently co-transfected with either A_1_R-Nluc or A_3_R-Nluc,β-arrestin1/2-YFP and hGRK1-6, or pcDNA3.1 vector (total 2 μg, at a AR:β-arrestin:hGRK ratio of 1:5:4) using polyethyleneimine (PEI, 1 mg/ml, MW = 25,000 g/mol; Polysciences Inc) at a DNA:PEI ratio of 1:6 (w/v). As a negative control for the A_1_R, transfections were also set up in the absence of β-arrestin1/2-YFP. Briefly, in sterile tubes containing 150 mM NaCl, DNA or PEI was added (final volume 50 μl) and allowed to incubate at room temperature for 5 minutes before mixing together and incubating for a further 10 minutes prior to adding the combined mix dropwise to the cells. 24 hours post-transfection, HEK 293T cell were harvested, resuspended in reduced serum media (MEM, NEAA; Thermo Fisher Scientific) supplemented with 1% L-glutamine (2 mM final; Thermo Fisher Scientific), 2% FBS and 1x antibiotic-antimycotic solution and seeded (50,000 cells/well) in a poly-L-lysine-coated (MW: 150,000-300,000 Da; Sigma-Aldrich) white 96-well plate (PerkinElmer Life Sciences). 24 hours post seeding, media was removed, cells gently washed in PBS and 90 μl of furimazine (4 μM)-containing solution added (PBS supplemented with 0.49 mM MgCl_2_, 0.9 mM CaCl_2_ and 0.1% BSA) to each well before incubating in the dark for 10 minutes. After incubation, 10 μl of agonist (NECA, CPA, adenosine, BnOCPA) was added (0.01 μM to 10 μM) and filtered light emissions measured at 450 nm and 530 nm every minute for 1 hour using a Mithras LB 940 (Berthold technology). Here, Nluc on the C-terminus of A_1_R or A_3_R acted as the BRET donor (luciferase oxidizing its substrate) and YFP acted as the fluorescent acceptor. Vehicle control (1% DMSO) was added to determine background emission, and data was corrected for baseline reading, vehicle and the response obtained in the absence of YFP-β-arrestin1/2, when appropriate.

#### Radioligand binding

Radioligand displacement assays were conducted using crude membrane preparations (100 μg protein per tube) acquired from homogenisation of CHO-K1-hA_1_R cells in ice-cold buffer (2 mM MgCl_2_, 20 mM HEPES, pH 7.4). The ability to displace binding of the A_1_R-selective antagonist radioligand, 1,3-[^3^H]-dipropyl-8-cyclopentylxanthine ([^3^H]-DPCPX) at a concentration (1 nM) around the Kd value (1.23 nM, as determined by saturation binding experiments) by increasing concentrations of NECA, adenosine, CPA, BnOCPA or HOCPA (0.1 nM – 10 μM) allowed the binding affinities (Ki) to be determined. Non-specific binding was determined in the presence of 10 μM DPCPX. Membrane incubations were conducted in Sterilin™ scintillation vials (Thermo Fisher Scientific; Wilmington, Massachusetts, USA) for 60 minutes at room temperature. Free radioligand was separated from bound radioligand by filtration through Whatman® glass microfiber GF/B 25 mm filters (Sigma-Aldrich). Each filter was then placed in a Sterilin™ scintillation vial and radioactivity determined by: addition of 4 mL of Ultima Gold XR liquid scintillant (PerkinElmer), overnight incubation at room temperature and the retained radioactivity determined using a Beckman Coulter LS 6500 Multi-purpose scintillation counter (Beckman Coulter Inc.; Indiana, USA).

#### NanoBRET ligand-binding studies

Real-time pharmacological interactions between ligands and receptors was quantitated using NanoBRET as described previously^8^. In brief, using N-terminally Nluc-tagged rA_1_R-, rA_2_AR- or rA_3_R-expressing HEK 293 cell lines, competition binding assays were conducted. In all antagonist assays CA200645, which acts as a fluorescent antagonist with a slow off-rate^9^, was used, with the exception of the rat A_3_R where the fluorescent compound was AV039^10^. The data was fitted with the ‘one-site – Ki model’ derived from the Cheng and Prusoff equation, built into Prism to determine affinity (pK_i_) values for all unlabelled agonists at all AR subtypes assayed. For the hA_1_R we also performed an agonist binding competition assay using NECA-TAMRA (Noel et al., unpublished). Here data was fitted with the ‘two-site Ki model’, build into Prism to determine high affinity and low affinity values for the unlabelled agonists. For all ARs, filtered light emission at 450 nm and > 610 nm (640-685 nm band pass filter) was measured using a Mithras LB 940 and the raw BRET ratio calculated by dividing the 610 nm emission with the 450 nm emission. The Nluc acts as the BRET donor (luciferase oxidizing its substrate) and CA200645/AV039/NECA-TAMRA acted as the fluorescent acceptor. CA200645 was used at 25 nM, as previously reported^11^, AV039 was used at 100 nM (Barkan et al. 2019) and NECA-TAMRA at its Kd of 15.2 μM (Noel et al., unpublished). BRET was measured following the addition of the Nluc substrate, Furimazine (0.1 μM). Nonspecific binding was determined using a high concentration of unlabelled antagonist, DPCPX for rA_1_R, ZM241385 for the rA_2_AR and MRS 1220 for rA_3_R.

#### TRUPATH G protein dissociation assay

Cells were plated in a density of 1,500,000 cells/well in a 6-well plate and grown in DMEM /F-12 GlutaMAX™ media (Thermo Fisher Scientific, UK) supplemented with 10% FBS (Sigma, UK) and 1% AA (Sigma, UK). After being grown overnight, cells in each well were transfected using polyethylenimine 25 kDa (PEI, Polysciences Inc., Germany) at a 6:1 ratio of PEI to DNA, diluted in 150mM NaCl. Cells were transfected with hA_1_R, Goa-RLuc8 or Gob-RLuc8, Gβ_3_, Gγ_8_-GFP2, and pcDNA3.1 with the ratio of 1:1:1:1:1 (400 ng per construct) in accordance with previously published methods^12^. Gɑ (either Goa-RLuc8, or Gob-RLuc8), Gβ3 and Gγ8-GFP2 constructs were purchased as part of the TRUPATH sensor kit from Addgene, pcDNA3.1-A_1_R was obtained from cDNA resource centre, and pcDNA3.1 (-) zeo was purchased from Invitrogen. After 24h, cells were trypsinised and re-seeded onto poly-L-lysine (PLL)-coated white 96-well plates (Greiner, UK) at the density of 50,000 cells/well in a complete DMEM/F12 medium. After grown overnight, the culture media was discarded and replaced with 80 μl assay buffer (1× Hank’s balanced salt solution (HBSS) with calcium, supplemented with 20 mM HEPES and 0.1% BSA at pH 7.4). The assay was started by adding 10 μl of coelenterazine 400a (Nanolight technology, USA) to a final concentration of 5 μM. The plates were then incubated in the dark for 5 minutes, prior to the addition of 10μl compounds (in a range of 0.01 nM – 1 µM). In order to investigate the effect of interfering peptides on Goa and Gob activation, cells were transfected with the TruPath constructs for Goa and Gob with the A_1_R as described above. However, the vector was replaced by either interfering or scrambled peptides, as appropriate, with increasing concentration: 0, 100, and 400 ng and was complemented by pcDNA3.1(-) up to 400 ng. CPA 10 µl was used as the ligand in a range of 1nM – 1 µM. BRET signal was recorded for 30 minutes on a Mithras LB940 plate reader allowing sequential integration of signal detected from GFP2 and Rluc8. The BRET ratio corresponds to the ratio of light emission from GFP2 (515 nm) over Rluc8 (400 nm). Net BRET ratio was used to generate the concentration response curve by taking 11-minute time-point after baseline correction. Data was analysed as change in the presence of the interfering peptides relative to control alone at 1 µM CPA.

#### Data Analysis

Concentration-response curves for the effects of A_1_R agonists on synaptic transmission were constructed in OriginPro 2018 (OriginLab; Northampton, MA, USA) and fitted with a logistic curve using the Levenberg Marquadt iteration algorithm. OriginPro 2018 was also used for statistical analysis. Statistical significance was tested as indicated in the text using paired or unpaired t-tests or one-way or two-way ANOVAs with repeated measures (RM) as appropriate. Bonferroni corrections for multiple comparisons were performed. All *in vitro* cell signalling assay data was analysed using Prism 8.4 (Graphpad software, San Diego, CA), with all concentration-response curves being fitted using a 3 parameter logistic equation to calculate response range and IC_50_. All cAMP data was normalised to a forskolin concentration-response curve ran in parallel to each assay. Where appropriate the operational model of receptor agonism^7, 13^ was used to obtain efficacy (log τ) and equilibrium disassociation constant (log *K_A_*) values. Calculation of bias factors (Δlog(Tau/K_A_)) relative to adenosine was performed as described in Weston *et al.* (2016)^7^. Error for this composite measure was propagated by applying the following equation.

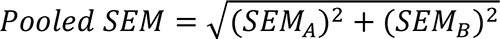

Where, σ_A_ and σ_B_ are the standard deviations of measurement A and B with mean of 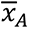 and 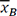 is the composite mean and n is the number of repeats.

Single-dose Schild analysis was performed on data using BnOCPA as an antagonist to adenosine in the cAMP assays so enabling determination of BnOCPA’s affinity constant (KA) using the following equation

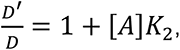

where D’ and D = EC_50_ values of adenosine with and without BnOCPA present, respectively, [A] = the concentration of BnOCPA, and K_2_ = the affinity constant (K_A_) of the BnOCPA^14^.

Statistical significance relative to adenosine was calculated using a one-way ANOVA with a Dunnett’s post-test for multiple comparisons. Radioligand displacement curves were fitted to the one-site competition binding equation yielding log(Ki) values. One-way ANOVA (Dunnett’s post-test) was used to determine significance by comparing the log(Ki) value for each compound when compared to adenosine. To determine the extent of ligand-induced recruitment of β-arrestin2-YFP to either the A_1_R or A_3_R, the BRET signal was calculated by subtracting the 530 nm/450 nm emission for vehicle-treated cells from ligand-treated cells (ligand-induced ΔBRET). ΔBRET for each concentration at 5 minutes (maximum response) was used to produce concentration-response curves.

All *in vivo* cardiovascular and respiratory data were analysed using OriginPro 2018. One-way ANOVAs, with repeated measures as appropriate, and with Bonferroni correction for multiple comparisons were used. Statistical significance for the effects of IV saline and the antagonist effect of BnOCPA on CPA were tested using paired t-tests. Data are reported throughout as mean ± SEM and n values are reported for each experiment. For the neuropathic pain studies, one-way ANOVAs with Fisher’s Least Significant Difference (LSD) post-hoc test was used to compare drug treatment groups to the vehicle group (OriginPro 2018). The significance level was set at P < 0.05, with actual P values reported in the figure legends and summaries, by way of abbreviations and asterisks, on the graphs: ns, not significant; * P < 0.05; **, P < 0.02; ***, P < 0.001; ****, P < 0.0001.

#### Drugs and substances

Drugs were made up as stock solutions (1-10 mM) and then diluted in aCSF or saline on the day of use. BnOCPA^15^ ((2*R*,3*R*,4*S*,5*R*)-2-(6-{[(1*R*,2*R*)-2-benzyloxycyclopentyl]amino}-9*H*-purin-9-yl)-5-(hydroxymethyl)oxolane-3,4-diol) and HOCPA^16^ ((2*R*,3*R*,4*S*,5*R*)-2-(6-{[(1*R*,2*R*)-2-hydroxycyclopentyl]amino}-9*H*-purin-9-yl)-5-(hydroxymethyl)oxolane-3,4-diol), the [(1*R*,2*R*)-2-hydroxycyclopentyl]amino bis-epimer of known A_1_R agonist GR79236^17^, were synthesised as described previously^6^ and dissolved in dimethyl-sulphoxide (DMSO, 0.01% final concentration). Adenosine, 8-CPT (8-cyclopentyltheophylline), NECA (5′-(*N*-Ethylcarboxamido) adenosine), DPCPX, ZM241385, MRS1220 and CPA (*N*^6^-Cyclopentyladenosine) were purchased from Sigma-Aldrich (Poole, Dorset, UK). MRS 1523 was purchased from Cayman Chemicals (Cambridge Bioscience Ltd., Cambridge UK). [^3^H]-DPCPX was purchased from PerkinElmer (Life and Analytical Sciences, Waltham, MA). CA200645 and peptides for interfering with G protein signalling were obtained from Hello Bio (Bristol, UK) and were based on published sequences^5^. NECA-TAMARA was synthesised in house (Noel *et al.*, in preparation), while AV039, a highly potent and selective fluorescent antagonist of the human A_3_R based on the 1,2,4-Triazolo[4,3-a]quinoxalin-1-one linked to BY630 was kindly gifted to us by Stephen Hill and Stephen Briddon (University of Nottingham). For Goa the peptide had a sequence of MGIANNLRGCGLY. The scrambled version was LNRGNAYLCIGMG. For Gob the peptide had a sequence of MGIAKNLRGCGLY. Peptides were made up as stock solutions (2 mM) and stored at – 20°C. The stock solutions were dissolved in filtered intracellular solution just before use.

#### BnOCPA Pharmacokinetics

The stability in solution and metabolism of BnOCPA (0.1 µM or 1 µM) was assessed by Eurofins Panlabs. The parameters examined were: half-life (t_1/2_) in PBS (1 µM BnOCPA, 37 °C, pH 7.4; Assay #600); t_1/2_ in human plasma (1 µM BnOCPA, 37 °C; Assay #887) and intrinsic clearance by human liver microsomes (0.1 µM BnOCPA, 0.1 mg/ml, 37 °C; Assay #607).

#### Half-life determination in PBS

At the end of the incubation at each of the time points (0, 1, 2, 3, 4 hours), an equal volume of an organic mixture (acetonitrile/methanol, 50/50 v/v) was added to the incubation mixture. Samples were analyzed by HPLC-MS/MS and corresponding peak areas were recorded for each analyte. The ratio of precursor compound remaining after each time point relative to the amount present at time 0, expressed as a percentage, is reported as chemical stability. The t_1/2_ was estimated from the slope of the initial linear range of the logarithmic curve of compound remaining (%) versus time, assuming first order kinetics.

#### Half-life determination in human plasma

At the end of incubation at each of the time points (0, 0.5, 1, 1.5, 2 hours), acetonitrile was added to the incubation mixture followed by centrifugation. Samples were analyzed by HPLC-MS/MS and peak areas were recorded for each analyte. The area of precursor compound remaining after each of the time points relative to the amount remaining at time 0, expressed as a percentage, was calculated. Subsequently, the t_1/2_ is estimated from the slope of the initial linear range of the logarithmic curve of compound remaining (%) versus time, assuming first order kinetics.

#### Intrinsic clearance by human liver microsomes

Metabolic stability, expressed as a percentage of the parent compound remaining, was calculated by comparing the peak area of the compound at the time point (0, 15, 30, 45, 60 minutes) relative to that at time 0. The t_1/2_ was estimated from the slope of the initial linear range of the logarithmic curve of compound remaining (%) vs. time, assuming the first-order kinetics. The apparent intrinsic clearance (CL_int_, in µL/min/mg) was calculated according to the following formula:

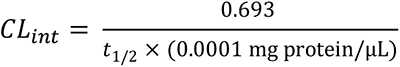

The behaviour of BnOCPA was compared to appropriate standards. Data is available in Supplemental Data File 1

### Molecular Dynamics Simulations

#### Ligand parameterization

The CHARMM36^18, 19^/CGenFF^20–22^ force field combination was employed in all the molecular dynamic (MD) simulations performed. Initial topology and parameter files of BnOCPA, HOCPA, and PSB36 were obtained from the Paramchem webserver^20^. Higher penalties were associated with a few BnOCPA dihedral terms, which were therefore optimized at the HF/6-31G* level of theory using both the high throughput molecular dynamics (HTMD)^23^ parameterize functionality and the Visual Molecular Dynamics (VMD)^24^ Force Field Toolkit (ffTK)^25^, after fragmentation of the molecule. Short MD simulations of BnOCPA in water were performed to visually inspect the behavior of the optimized rotatable bonds.

#### Systems preparation for fully dynamic docking of BnOCPA and HOCPA

Coordinates of the A_1_R in the active, adenosine- and G protein-bound state were retrieved from the Protein Data Bank^26, 27^ database (PDB ID 6D9H^28^). Intracellular loop 3 (ICL3) which is missing from PDB ID 6D9H was rebuilt using Modeller 9.19^29, 30^. The G protein, with the exception of the C-terminal helix (helix 5) of the G protein alpha subunit (the key region responsible for the receptor TM6 active-like conformation) was removed from the system as in previous work^31, 32^. BnOCPA and HOCPA were placed in the extracellular bulk, in two different systems, at least 20 Å from the receptor vestibule. The resulting systems were prepared for simulations using in-house scripts able to exploit both python HTMD^23^ and Tool Command Language (TCL) scripts. Briefly, this multistep procedure performs the preliminary hydrogen atoms addition by means of the pdb2pqr^33^ and propka^34^ software, considering a simulated pH of 7.0 (the proposed protonation of titratable side chains was checked by visual inspection). Receptors were then embedded in a square 80 Å x 80 Å 1-palmitoyl-2-oleyl-sn-glycerol-3-phosphocholine (POPC) bilayer (previously built by using the VMD Membrane Builder plugin 1.1, Membrane Plugin, Version 1.1.; http://www.ks.uiuc.edu/Research/vmd/plugins/membrane/) through an insertion method^35^, considering the A_1_R coordinates retrieved from the OPM database^36^ to gain the correct orientation within the membrane. Lipids overlapping the receptor transmembrane bundle were removed and TIP3P water molecules^37^ were added to the simulation box (final dimensions 80 Å × 80 Å × 125 Å) using the VMD Solvate plugin 1.5 (Solvate Plugin, Version 1.5; http://www.ks.uiuc.edu/Research/vmd/plugins/solvate/). Finally, overall charge neutrality was achieved by adding Na^+^/Cl^-^ counter ions (concentration of 0.150 M) using the VMD Autoionize plugin 1.3 (Autoionize Plugin, Version 1.3; http://www.ks.uiuc.edu/Research/vmd/plugins/autoionize/). All histidine side chains were considered in the delta tautomeric state, with the exception of H251 (epsilon tautomer) and H278 (protonated).

The MD engine ACEMD^38^ was employed for both the equilibration and productive simulations. Systems were equilibrated in isothermal-isobaric conditions (NPT) using the Berendsen barostat^39^ (target pressure 1 atm), the Langevin thermostat^40^ (target temperature 300 K) with a low damping factor of 1 ps^-1^ and with an integration time step of 2 fs. Clashes between protein and lipid atoms were reduced through 2000 conjugate-gradient minimization steps before a 2 ns long MD simulation was run with a positional constraint of 1 kcal mol^-1^ Å^-2^ on protein and lipid phosphorus atoms. Twenty nanoseconds of MD simulation were then performed constraining only the protein atoms. Lastly, positional constraints were applied only to the protein backbone alpha carbons for a further 5 ns.

#### Dynamic docking of BnOCPA and HOCPA

The supervised MD (SuMD) approach is an adaptive sampling method^41^ for simulating binding events in a timescale one or two orders of magnitudes faster than the corresponding classical (unsupervised) MD simulations^42^. SuMD has been successfully applied to small molecules and peptides^43–49^. In the present work, the distances between the centers of mass of the adenine scaffold of the A_1_R agonist and N254^6.55^, F171^ECL2^, T277^7.42^ and H278^7.43^ of the receptor were considered for the supervision during the MD simulations. The dynamic docking of BnOCPA was hindered by the ionic bridge formed between the E172^ECL2^ and K265^ECL3^ side chains. A metadynamics^50–52^ energetic bias was therefore introduced in order to facilitate the rupture of this ionic interaction, thus favoring the formation of a bound complex. More precisely, Gaussian terms (height = 0.01 kcal mol^-1^ and widths = 0.1 Å) were deposited every 1 ps along the distance between the E172^ECL2^ carboxyl carbon and the positively charged K265^ECL3^ nitrogen atom using PLUMED 2.3^53^. A similar SuMD-metadynamics hybrid approach was previously employed to study binding/unbinding kinetics^54^ on the A_2_AR subtype. For each replica (Methods Table 1), when the ligands reached a bound pose (i.e. a distance between the adenine and the receptor residues centers of mass < 3 Å), a classic (unsupervised and without energetic bias) MD simulation was performed for at least a further 100 ns.

#### BnOCPA bound state metadynamics

We decided to perform a detailed analysis of the role played by the E172^ECL2^ – K265^ECL3^ ionic interaction in the dynamic docking of BnOCPA. Three 250 ns long well-tempered^55^ metadynamics simulations were performed using the bound state obtained from a previous dynamic docking simulation, which resulted in binding mode A, as a starting point. The collective variables (CVs) considered were: i) the distance between the E172^ECL2^ carboxyl carbon and the positively charged K265^ECL3^ nitrogen atom and ii) the dihedral angle formed by the 4 atoms linking the cyclopentyl ring to the phenyl moiety (which was the most flexible ligand torsion during the previous SuMD simulations). Gaussian widths were set at 0.1 Å and 0.01 radians respectively, heights at 0.01 kcal/mol^-1^, and the deposition was performed every 1 ps (bias-factor = 5). Although complete convergence was probably not reached, three replicas (Methods Table 1) allowed sampling of three main energetic minima on the energy surface (Supplementary Fig. 8); these correspond to the representative binding poses shown in Fig. 3d to f.

#### Classic MD simulations of BnOCPA binding modes A, B, C and D

To test the hypothesis that BnOCPA and HOCPA may differently affect TM6 and/or TM7 mobility when bound to A_1_R (and to further sample the stability of each BnOCPA binding mode), putative binding conformations A, B and C (Fig. 3) were superposed to the experimental A_1_R active state coordinates with the modelled ICL3. This should have removed any A_1_R structural artefacts, possibly introduced by metadynamics. As reference and control, two further systems were considered: i) the pseudo-apo A_1_R and ii) the selective A_1_R antagonist PSB36^56^ superposed in the same receptor active conformation (Methods Table 1). The BnOCPA binding mode D was modelled from mode B by rotating the dihedral angle connecting the cyclopentyl ring and the N6 nitrogen atom in order to point the benzyl of the agonist toward the hydrophobic pocket underneath ECL3 (Fig. 3g) delimited by L253^6.56^, T257^6.52^, K265^ECL3^, T270^7.35^, and L269^7.34^. The G protein atoms were removed, and the resulting systems prepared for MD as reported above. A similar comparison was performed in a milestone study on the β_2_ adrenergic receptor^57^ which sought to describe the putative deactivation mechanism of the receptor.

#### Dynamic docking of the Goa, Gob and Gi2 GαCT helix

A randomly extracted frame from the classic MD performed on the BnOCPA:A_1_R complex was prepared for three sets of simulations placing the GαCT helix 5 (last 27 residues) of the Gα proteins Goa, Gob and Gi2 in the intracellular solvent bulk side of the simulation boxes. As a further control, a frame from the classic MD performed on the unbiased ligand HOCPA:A_1_R complex was randomly extracted and prepared along with the Gob GαCT. The resulting four systems were embedded in a POPC membrane and prepared as reported above.

The different structural effects putatively triggered by BnOCPA and HOCPA on the recognition mechanism of Goa, Gob and Gi2 GαCT were studied by performing 10 SuMD replicas (Methods Table 1). During each replica (Video S3), the distance between the centroid of the GαCT residues 348-352 and the centroid of the A_1_R residues D42^2.37^, I232^6.33^, and Q293^8.48^ was supervised until it reached a value lower than 8 Å. A classic MD simulation was then run for a further 300 ns.

#### Classic MD simulations on the A_1_R:Goa and Gob complexes

The A_1_R cryo-EM structure (PDB ID 6D9H) was used as template for all the five systems simulated (Methods Table 1). The endogenous agonist adenosine was removed and HOCPA and BnOCPA (modes B and D) were inserted in the orthosteric site superimposing 6D9H to the systems prepared for the classic MD simulations in the absence of G protein. ICL3 was not modelled, nor were the missing part of the G protein α subunit. As subunits β and γ were removed, the Gα NT helix was truncated to residue 27 to avoid unnatural movements (NT is constrained by Gβ in 6D9H). The Gα subunit was mutated according to the Goa and Gob primary sequences^58^ using in-house scripts. The resulting five systems (Methods Table 1) were embedded in a POPC membrane and prepared as reported above.

#### Analysis of the classic MD simulations

During the classic MD simulations that started from Modes A-C (Fig. 3d to f), BnOCPA had the tendency to explore the three conformations by rapidly interchanging between the three binding modes. In order to determine the effect exerted on the TM domain by each conformation, 21 μs of MD simulations (Methods Table 1 – BnOCPA mode A, BnOCPA mode B, BnOCPA mode C) were subjected to a geometric clustering. More precisely, a simulation frame was considered in pose A if the distance between the phenyl ring of BnOCPA and the I175^ECL2^ alpha carbon was less than 5 Å; in pose B if the distance between the phenyl ring of BnOCPA and the L258^6.59^ alpha carbon was less than 6 Å, and in pose C if the distance between the phenyl ring of BnOCPA and the Y271^7.36^ alpha carbon was less than 6 Å. During the MD simulations started from mode D (Fig. 3g), a frame was still considered in mode D if the root mean square deviation (RMSD) of the benzyl ring to the starting equilibrated conformation was less than 3 Å. For each of the resulting four clusters, the RMSD of the GPCR conserved motif NPXXY (N^7.49^ PIV Y^7.53^ in the A_1_R; Supplementary Fig. 9) was computed using Plumed 2.3^53^ considering the inactive receptor state as reference, plotting the obtained values as frequency distributions (Fig. 3i, Rearrangement of the NPXXY motif, which is located at the intracellular half of TM7, is considered one of the structural hallmarks of GPCR activation^59^. Upon G protein binding, it moves towards the center of the receptor TM bundle (Supplementary Fig. 9). Unlike other activation micro-switches (e.g. the break/formation of the salt bridge between R^3.50^ and E^6.30^), this conformational transition is believed to occur in timescales accessible to MD simulations^57^.

Hydrogen bonds and atomic contacts were computed using the GetContacts analysis tool (https://github.com/getcontacts/getcontacts) and expressed in terms of occupancy (the percentage of MD frames in which the interaction occurred).

#### Analysis of the Goa, Gob and Gi2 GαCT classic MD simulations after SuMD

For each system, only the classic MD simulations performed after the GαCT reached the A_1_R intracellular binding site were considered for the analysis.

The RMSD values to the last 15 residues of the Gi2 GαCT reported in the A_1_R cryo-EM PDB structure 6D9H were computed using VMD^24^. The MD frames associated with the peaks in the RMSD plots (states CS1, MS1, MS2 and MS3 in Fig. 4a, d) were clustered employing the VMD Clustering plugin (https://github.com/luisico/clustering) by selecting the whole GαCT helixes alpha carbon atoms and a cutoff of 3 Å.

**Methods Table 1.** Summary of the simulations performed.

| Ligand | MD approach | # Replicas | Total simulated time<sup>a</sup> |
| --- | --- | --- | --- |
| BnOCPA | SuMD | 6 | 1.9 μs |
| BnOCPA | SuMD-Metadynamics | 5 | 4.3 μs |
| HOCPA | SuMD | 5 | 3.4 μs |
| BnOCPA (bound state after dynamic docking) | Metadynamics | 3 | 0.75 μs |
| BnOCPA(A) | Classic MD | 6 | 9.0 μs |
| BnOCPA(B) | Classic MD | 6 | 9.0 μs |
| BnOCPA(C) | Classic MD | 3 | 3.0 μs |
| HOCPA | Classic MD | 4 | 8.0 μs |
| PSB36 | Classic MD | 4 | 6.0 μs |
| Apo A<sub>1</sub> | Classic MD | 4 | 8.0 μs |
| GαCT Goa (BnOCPA) | SuMD + Classic MD | 10 | 0.36 μs + 3.0 μs |
| GαCT Gob (BnOCPA) | SuMD + Classic MD | 10 | 0.33 μs + 3.0 μs |
| GαCT Gi2 (BnOCPA) | SuMD + Classic MD | 10 | 0.37 μs + 3.0 μs |
| GαCT Gob (HOCPA) | SuMD + Classic MD | 10 | 0.29 μs + 3.0 μs |
| BnOCPA(D):Gob | Classic MD | 4 | 4.0 μs |
| BnOCPA(B):Gob | Classic MD | 3 | 3.0 μs |
| HOCPA:Gob | Classic MD | 4 | 4.0 μs |
| BnOCPA(D):Goa | Classic MD | 5 | 5.0 μs |
| BnOCPA(B):Goa | Classic MD | 4 | 4.0 μs |

a) For SuMD and SuMD-metadynamics simulations the time is the sum of productive SuMD time windows.
(A), (B), (C) and (D) indicate the respective BnOCPA binding modes.

